# Translational adaptation to heat stress is mediated by 5-methylcytosine RNA modification in *Caenorhabditis elegans*

**DOI:** 10.1101/2020.03.21.001735

**Authors:** Isabela Cunha Navarro, Francesca Tuorto, David Jordan, Carine Legrand, Jonathan Price, Fabian Braukmann, Alan Hendrick, Alper Akay, Annika Kotter, Mark Helm, Frank Lyko, Eric A. Miska

## Abstract

Methylation of carbon-5 of cytosines (m^5^C) is a post-transcriptional nucleotide modification of RNA found in all kingdoms of life. While individual m^5^C-methyltransferases have been studied, the impact of the global cytosine-5 methylome on development, homeostasis and stress remains unknown. Here, using *Caenorhabditis elegans*, we generated the first organism devoid of m^5^C in RNA, demonstrating that this modification is non-essential. We determined the localisation and enzymatic specificity of m^5^C sites in RNA *in vivo* and showed that animals devoid of m^5^C are sensitive to temperature stress. At the molecular level, we showed that loss of m^5^C specifically impacts decoding of leucine and proline thus reducing the translation efficiency of transcripts enriched in these amino acids. Finally, we found translation of leucine UUG codons to be the most strongly affected upon heat shock, suggesting a role of m^5^C tRNA wobble methylation in the adaptation to heat stress.

## INTRODUCTION

First described in 1958 (Amos & Korn, 1958), the methylation of carbon-5 of cytosines (m^5^C) in RNAs is now known to be a conserved modification in biological systems. m^5^C has been detected in tRNAs, rRNAs, mRNAs and non-coding RNAs, and is catalyzed by m^5^C RNA-methyltransferases that are encoded in genomes from bacteria to humans (Boccaletto et al., 2017). In humans, m^5^C formation in RNA is catalysed by the tRNA aspartic acid MTase 1 (TRDMT1), also referred as DNMT2, as well as by members of the NOP2/Sun domain family of proteins, which contains seven members (NSUN1-7) (reviewed in (García-Vílchez, Sevilla, & Blanco, 2019).

In the last few years m^5^C has been implicated in a variety of molecular roles, according to its presence in different RNA classes. Reports published thus far showed that m^5^C methylation of tRNAs is catalyzed by NSUN2, NSUN3, NSUN6 and DNMT2. NSUN2 mediates the methylation of positions C48-50 in several cytoplasmic tRNAs and position C34 of tRNA Leu-CAA (Blanco et al., 2014; Brzezicha et al., 2006; Khoddami & Cairns, 2013; Martinez et al., 2012; Motorin & Grosjean, 1999; Squires et al., 2012; Tuorto et al., 2012). Additionally, a recent report has demonstrated that NSUN2 is imported into the matrix of mammalian mitochondria, where it catalyses m^5^C methylation of positions 48-50 in several mitochondrial tRNAs (Van Haute et al., 2019). Loss of NSUN2-mediated tRNA methylation has been shown to promote cleavage by angiogenin and accumulation of tRNA fragments that interfere with the translation machinery (Blanco et al., 2016, 2014; Flores et al., 2016). Despite using a DNA methyltransferase-like catalytic mechanism (Jurkowski et al., 2008), DNMT2 possesses targeting specificity towards position C38 of tRNAs Asp, Gly and Val (Goll et al., 2006; Schaefer et al., 2010). Similarly to NSUN2, DNMT2-mediated methylation was found important for protection of tRNAs from stress-induced cleavage in *Drosophila* and mice (Schaefer et al., 2010; Tuorto et al., 2015). At the translational level, it has been shown to facilitate charging of tRNA Asp and discrimination between Asp and Glu near-cognate codons, thus controlling the synthesis of Asp-rich sequences and promoting translational fidelity (Shanmugam et al., 2015; Tuorto et al., 2015). The m^5^C methyltransferase NSUN3 methylates exclusively mitochondrial tRNA Met-CAU at the wobble position (C34). This position is further modified into f^5^C by the dioxygenase ALKBH1 (Haag et al., 2016; Nakano et al., 2016; Van Haute et al., 2016), and facilitates the recognition of AUA and AUG codons as methionine in the mitochondria (Takemoto et al., 2009). Lack of tRNA Met-CAU C34 modification has been shown to affect mitochondrial translation rates in human fibroblasts (Van Haute et al., 2016). NSUN6 shows targeting specificity towards position C72 at the 3′-end of the acceptor stem of tRNAs Cys and Thr (Haag et al., 2015). Functionally, m^5^C methylation at position 72 was shown to promote a slight enhancement of thermal stability in tRNAs (J. Li et al., 2018).

m^5^C methylation in rRNA is mediated by NSUN1, NSUN4 and NSUN5. NSUN1 (yeast *nop2*) methylates C2870 in yeast 25S rRNA. While NSUN1 is an essential gene in all organisms studied thus far, reports focusing on the yeast homologue provided contradictory results regarding the phenotypes observed in organisms lacking its catalytic activity (Bourgeois et al., 2015; Sharma, Yang, Watzinger, Kötter, & Entian, 2013). A similar phenomenon has been observed for NSUN4, a mitochondrial rRNA m^5^C methyltransferase with targeting capability towards C911 of 12S mt-rRNA in mice (Cámara et al., 2011; Metodiev et al., 2014; Spåhr, Habermann, Gustafsson, Larsson, & Hallberg, 2012; Yakubovskaya et al., 2012). NSUN4 acts in complex with MTERF4 for assembly of small and large mitochondrial ribosome subunits, however m^5^C catalysis occurs in an MTERF4-independent manner and does not seem to be essential (Metodiev et al., 2014). NSUN5 has been shown to methylate position C2278 in yeast 25S rRNA (Schosserer et al., 2015; Sharma et al., 2013). Loss of NSUN5-mediated methylation induces conformational changes in the ribosome and modulation of translational fidelity, favouring the recruitment of stress-responsive mRNAs into polysomes and promoting lifespan enhancement in yeast, flies and nematodes (Schosserer et al., 2015). In mice, Nsun5 knockout causes reduced body weight and reduced protein synthesis in several tissues (Heissenberger et al., 2019). Finally, NSUN7 has been shown to methylate eRNAs associated to targets of the metabolic activator PGC-1α, enhancing eRNA stability and upregulating PGC-1α target genes (Aguilo et al., 2016).

Several methods have been used to investigate the presence of m^5^C in coding transcripts. However, a general consensus on the abundance, distribution and relevance of this mark in mRNAs has not yet been achieved, as the number of putative mRNA m^5^C sites varies up to 1,000-fold between studies (David et al., 2017; Huang, Chen, Liu, Gu, & Zhang, 2019; Legrand et al., 2017; Q. Li et al., 2017; Squires et al., 2012; Tang et al., 2015; Yang et al., 2017; Zhang et al., 2012). Recently, further mechanistic insights into potential functions of m^5^C in coding transcripts have been proposed through the identification of the reader proteins ALYREF and YBX1. These proteins have been shown to bind m^5^C sites in mRNAs to modulate nuclear-cytoplasm transport and mRNA stability, respectively (Chen et al., 2019; Yang et al., 2017). Pathogenic mutations in humans have been mapped to genes involved in the m^5^C pathway, indicative of the relevance of studying the basic molecular features behind this modification (Abbasi-Moheb et al., 2012; Haag et al., 2016; Khan et al., 2012; Khosronezhad, Golam, & Jorsarayi, 2015; Khosronezhad, Hosseinzadeh Colagar, & Mortazavi, 2015; Komara, Al-Shamsi, Ben-Salem, Ali, & Al-Gazali, 2015; Martinez et al., 2012; Nakano et al., 2016; Ren, Zhong, Ding, Chen, & Jing, 2015; Van Haute et al., 2016). Although previous studies have explored the roles of m^5^C methyltransferases individually, none of them has focused on establishing a simultaneous systematic description of these enzymes, of their specificity, or of their potential molecular and genetic interactions, in any organism. While it is clear that m^5^C can exert distinct roles in a wide variety of molecular contexts, it remains unclear how the m^5^C methylome acts as a whole to sustain development. Here we used *Caenorhabditis elegans* as a model to study the genetic requirements and molecular functions of the m^5^C pathway in a whole organism. We produced the first animal strain devoid of detectable levels of this modification and used this genetic tool to determine the localisation of m^5^C sites and their impact on physiology and stress.

## RESULTS

### m^5^C and its derivatives are non-essential RNA modifications in *C. elegans*

To identify potential m^5^C RNA methyltransferases in *C. elegans,* we performed a BLAST analysis and found that the open reading frames *W07E6.1*, *Y48G8AL.5*, *Y39G10AR.21* and *Y53F4B.4* are putative homologues of the human genes NSUN1, NSUN2, NSUN4 and NSUN5, respectively (**Figure 1A**). Knockdown of these genes through RNAi by feeding revealed that *nsun-1* is an essential gene, as 100% of the animals that had this gene silenced from the first larval stage developed into sterile adults (**Supplementary Figure 1A-B**). We could not find homologous genes of of NSUN3, NSUN6, NSUN7 or DNMT2. m^5^C RNA methyltransferases utilise two conserved cysteine residues for the methyl group transfer, one of which (TC-Cys) is required for the covalent adduct formation, and the other (PC-Cys) for the release of the substrate following m^5^C catalysis (King & Redman, 2002). Using CRISPR-Cas9, we introduced mutations converting the TC-Cys into alanine in *nsun-1* (*mj473*), *nsun-2* (*mj458*) and *nsun-4* (*mj457*). These mutants, as well as a previously reported knockout mutant of *nsun-5* (*tm3898*), are viable and produce viable progeny, suggesting that the individual activity of m^5^C methyltransferases is not essential for the viability of *C. elegans* (**Figure 1B, Supplementary Figure 1C-D**). In addition, these results suggest that the essential role played by NSUN-1 in fertility (**Supplementary Figure 1A-B**) relies on non-catalytic functions of this protein.

**Figure 1.**
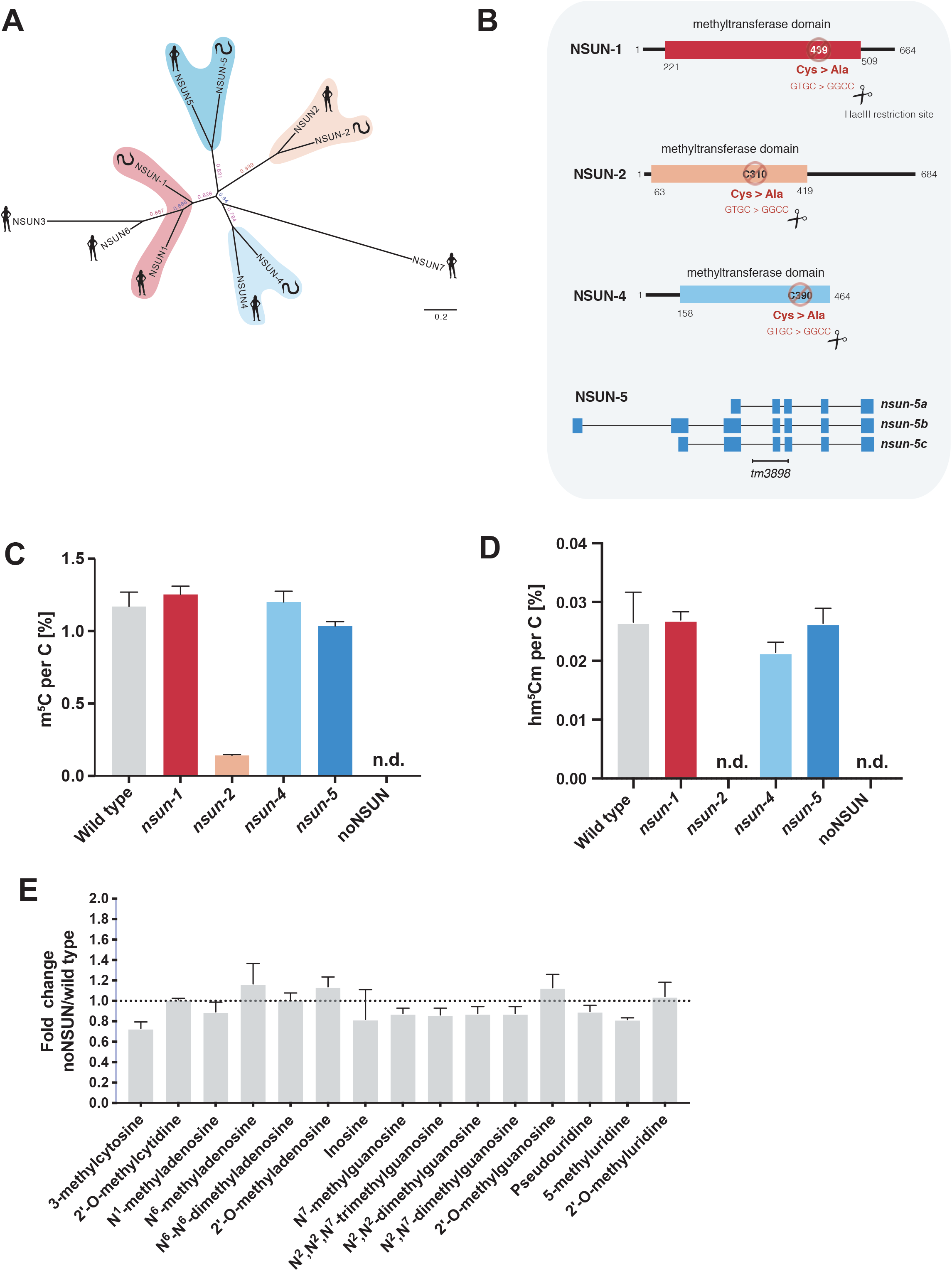
m^5^C and its derivatives are non-essential RNA modifications in *C. elegans*. **(A)** Phylogenetic relationship among human and putative nematode NSUN proteins. Unrooted phylogenetic tree of NSUN homologues in *Homo sapiens* and *C. elegans* using entire protein sequence. Different colours represent each member of the NSUN family: NSUN-1 (red); NSUN-2 (orange); NSUN-4 (light blue); NSUN-5 (dark blue). Phylogenetic tree reconstructed using the maximum likelihood method implemented in the PhyML program (v3.0). **(B)** Mutant alleles used in this study. CRISPR-Cas9 strategy for creation and screening of catalytically inactive alleles of *nsun-1* (*mj473*), *nsun-2* (*mj458*) and *nsun-4* (*mj457*). A homologous recombination template bearing a point mutation to convert the catalytic cysteine into alanine while creating a restriction site for HaeIII was co-injected. For the study of *nsun-5*, a 928 bp deletion allele (*tm3898*) was used. **(C, D)** Mass spectrometry quantification of m^5^C (C) and hm^5^Cm (D) levels in total RNA from *nsun* mutants. RNA was extracted from populations of L1 animals synchronised by starvation, digested to nucleosides and analysed via LC-MS. A relative quantification was performed with an external calibration curve of standards, normalized to the amount of cytidine. 3 independent biological replicates, mean with SEM. n.d. = not detected. **(E)** Fold change in total RNA modification levels upon loss of m^5^C. Fold changes were calculated by dividing the peak area ratio of noNSUN samples by the one of wild type samples. The dotted line represents the level where no change in RNA modification levels was observed. 3 independent biological replicates, mean with SEM; multiple t-tests.

To investigate epistatic interactions among *nsun* genes in *C. elegans*, we performed genetic crosses between the individual mutants, and produced a quadruple mutant in which all *nsun* genes are predicted to be inactive. This strain was viable and fertile, and was termed noNSUN. To confirm whether the introduced mutations resulted in catalytic inactivation, and to rule out the existence of additional unknown m^5^C RNA methyltransferases, we performed mass spectrometry analyses in total RNA from the mutant strains, and confirmed that m^5^C is no longer detectable in this genetic background (**Figure 1C**; lower limit of detection 316 pg/ml). We additionally quantified the m^5^C metabolic derivative 2′-O-methyl-5-hydroxymethylcytosine (hm^5^Cm) and found that this modification is not present either in *nsun-2* or noNSUN mutants (**Figure 1D**). We therefore conclude that m^5^C and its derivatives are not essential for *C. elegans* viability under laboratory conditions. Furthermore, we showed that NSUN-2 is the main source of m^5^C (88% of total levels), and that hm^5^Cm sites exclusively derive from NSUN-2 targets in *C. elegans*.

It has been proposed that some RNA modifications may act in a combinatorial manner, providing compensatory effects to each other (Hopper & Phizicky, 2003). This prompted us to investigate whether complete loss of m^5^C would significantly interfere with the levels of other RNA modifications. We performed a mass spectrometry analysis to quantify 15 different modifications in total RNA and found no significant differences between wild type and noNSUN samples (**Figure 1E**). While we acknowledge the sensitivity limitations of such a measurement performed in total RNA, taken together, the undetectable levels of m^5^C and lack of obvious changes to other modifications analysed establish the noNSUN strain as a highly specific genetic tool for the study of m^5^C distribution and function *in vivo*.

### The m^5^C methylome of *C. elegans*

Schosserer et al. demonstrated that position C2381 of 26S rRNA is methylated at carbon-5 by NSUN-5 in *C. elegans*, being involved in lifespan modulation (Schosserer et al., 2015). Nevertheless, the m^5^C methylome of this organism remained to be determined. We therefore used the noNSUN strain as a negative control for whole-transcriptome bisulfite sequencing (WTBS) analysis (Legrand et al., 2017), aiming to determine the localisation of m^5^C sites in *C. elegans* RNA at single nucleotide resolution.

We identified C5 methylation at positions C2982 and C2381 of 26S cytoplasmic rRNA and positions C628 and C632 of 18S mitochondrial rRNA (**Figure 2A**). Using alignment to rRNA of different organisms we found that position C2982 is a conserved NSUN1 target (Sharma et al., 2013), while C2381, as previously reported, is a conserved NSUN5 target (Schosserer et al., 2015; Sharma et al., 2013). We further confirmed the specificity of these sites using a targeted bisulfite sequencing (BS-seq) approach in individual mutants (**Supplementary Figure 2A, B**). Interestingly, other groups had previously identified adjacent modified sites in mt-rRNA in mice, however the methylation of only one of the positions was shown to be dependent on NSUN4 activity, while the other was interpreted as a 4-methylcytosine site (Metodiev et al., 2014). In the case of *C. elegans*, both positions are NSUN-dependent (**Figure 2A**). In addition, we could not detect m^4^C in wild type *C. elegans* total RNA by mass spectrometry, suggesting the occurrence of m^5^C in both positions in this organism (data not shown).

**Figure 2.**
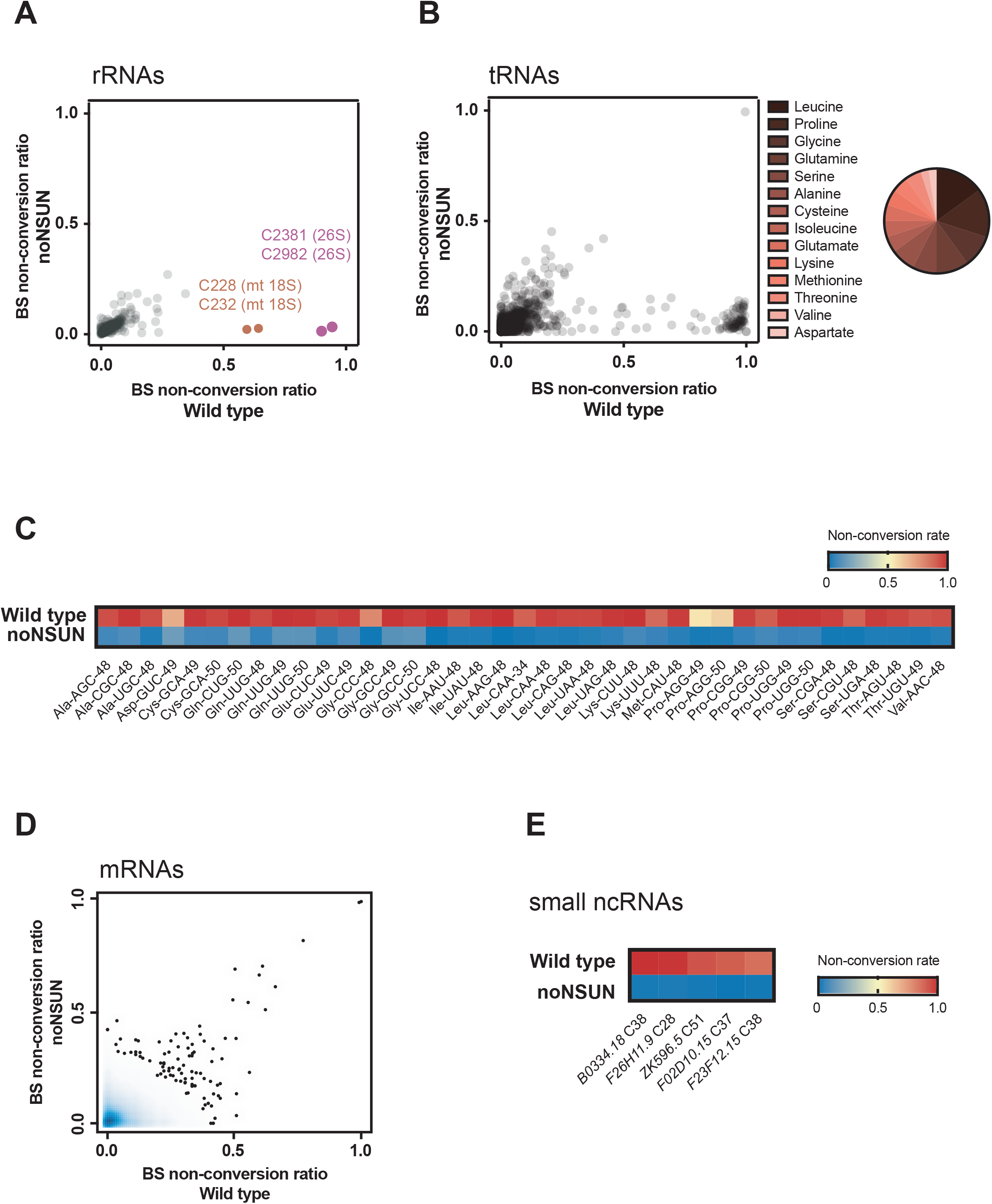
The m^5^C methylome of *C. elegans*. **(A, B)** Site-specific methylation analysis by whole-transcriptome bisulfite sequencing. Scatter plots show individual cytosines and their respective non-conversion ratios in rRNAs (A) and tRNAs (B) of wild type and noNSUN strains; pie chart showing most frequently methylated tRNA isoacceptors. **(C)** Heatmap showing non-conversion rates of tRNA positions methylated in stoichiometry higher than 50%. **(D)** Site-specific methylation analysis by whole-transcriptome bisulfite sequencing. Scatter plot shows density of cytosines and their respective non-conversion ratios in mRNAs of wild type and noNSUN strains. **(E)** Heatmap showing non-conversion rates of small non-coding RNA positions methylated in stoichiometry higher than 50%. 3 independent biological replicates.

We found 40 positions to be methylated in stoichiometry higher than 50% in tRNAs, the majority of which being detected in leucine and proline isoacceptors (**Figure 2B-C**). As anticipated for NSUN2 targeting, modified positions are found in the variable loop region (positions 48, 49, 50), with cytoplasmic tRNA Leu-CAA carrying an additional modification at the wobble position (C34) (Blanco et al., 2014; Burgess, David, & Searle, 2015). Using targeted BS-seq, we demonstrated that NSUN-2 is indeed responsible for both C34 and C48 methylation in tRNA-Leu (**Supplementary Figure 2C**). Surprisingly, we detected a position showing very high non-conversion ratio in an NSUN-independent manner (**Figure 2B**). In this case, the sequenced reads are identical to a 3′-fragment of tRNA Leu-CAA-2-1, apart from a C-T single nucleotide polymorphism. We predict that converted reads (bearing a T) would align to tRNA Leu-CAA-1-1, while non-converted (bearing a C) would align to tRNA Leu-CAA-2-1. Therefore, we treated this position as a probable alignment artefact.

Contrasting with what was observed for tRNAs and rRNAs, methylation of mRNAs in *C. elegans* is rare and in much lower stoichiometry. Lowering the non-conversion threshold from 50% to 25-40%, we detected 188 positions that remained unconverted after bisulfite treatment exclusively in wild type samples, *i.e.* putative m^5^C sites (**Figure 2D**, x axis). Using the same thresholds to probe for likely artefacts revealed 88 positions that remained unconverted exclusively in noNSUN samples, *i.e.* non-conversion was only 46% less frequent (**Figure 2D**, y axis). It is also noteworthy that positions remaining reproducibly highly unconverted equally in wild type and noNSUN samples are more frequent in mRNAs, when compared to tRNAs and rRNAs (**Figure 2A-B, D**). In summary, we found no evidence of a widespread distribution of m^5^C in coding transcripts. While our data does not completely rule out the existence of m^5^C methylation in mRNAs, it certainly demonstrates that this mark cannot be detected in high stoichiometry in *C. elegans*, as observed in tRNAs and rRNAs.

Finally, we attempted to identify m^5^C sites in small RNAs in our dataset. Given that the fractionation used in this protocol aimed to enrich for tRNAs (60-80 nt), we could not detect microRNA reads in abundance for confident analysis. Nevertheless, we found high NSUN-dependent non-conversion rates (>80%) in five non-coding RNAs (approximately 60 nt long) previously identified in *C. elegans* (Lu et al., 2011; Xiao et al., 2012). Secondary structure predictions suggest that the methylated sites are often present in a “hinge-like” context, or on the base of a stem-loop (**Supplementary Figure 3**). Further experiments will be required to determine the functionality of these m^5^C sites.

### NSUN-4 is a multisite-specific tRNA/rRNA-methyltransferase in the mitochondria of *C. elegans*

Interestingly, we also found the mitochondrial tRNA Met-CAU to be methylated at a very high rate (94.7%) (**Figure 3A**). Supporting our findings, previous articles also indicated the detection of this modified site in *Ascaris suum* and *C. elegans* (Nakano et al., 2016; Watanabe et al., 1994). This was unexpected, as previous reports have shown that this position is methylated by NSUN3 (Haag et al., 2016; Nakano et al., 2016; Van Haute et al., 2016). As *C. elegans* does not have an NSUN3 homologue, this implies that mitochondrial tRNAs can be modified by alternative enzymes.

**Figure 3.**
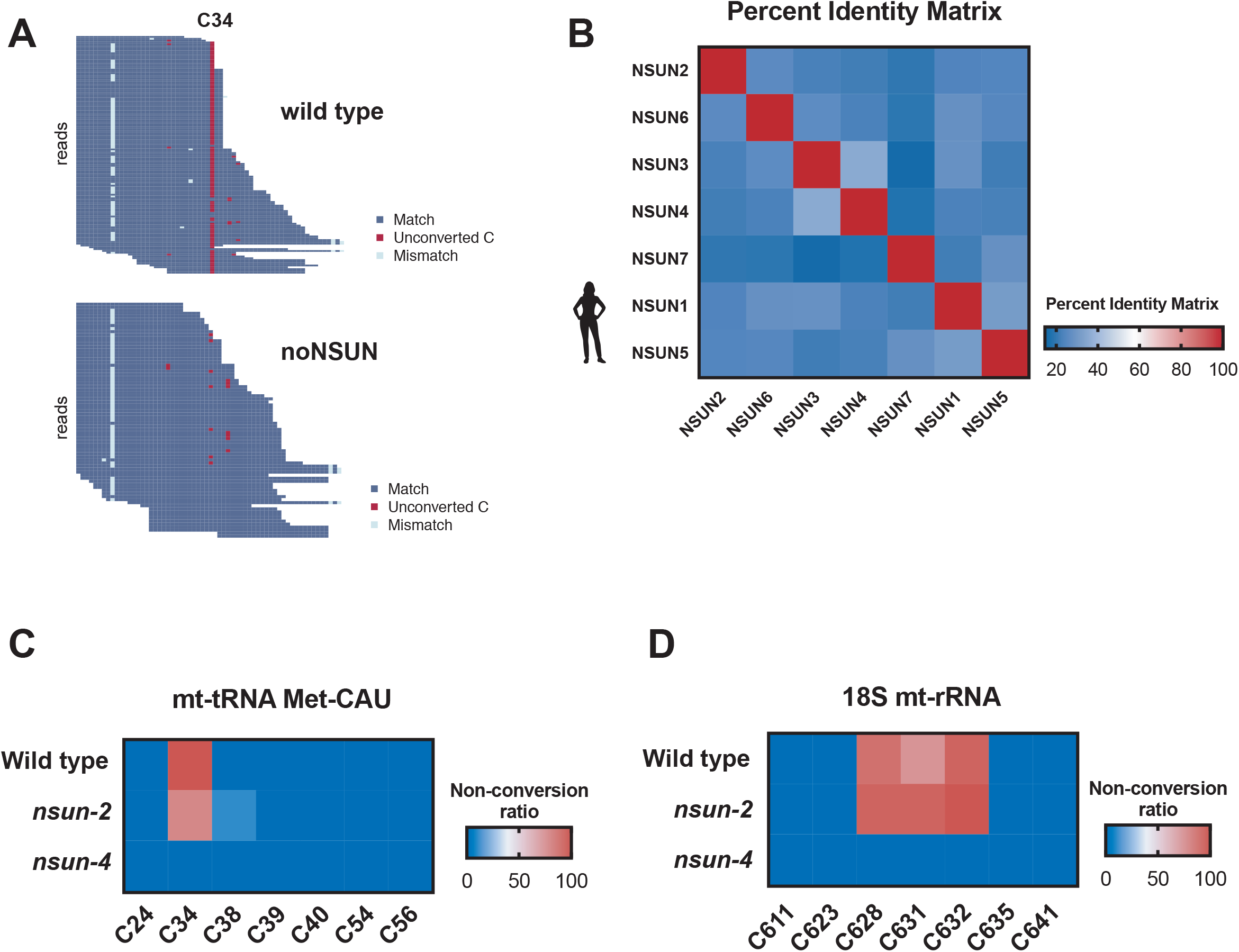
NSUN-4 is a multisite-specific tRNA/rRNA methyltransferase in *C. elegans*. **(A)** RNA bisulfite sequencing map for mitochondrial tRNA Met-CAU in wild type (top) and noNSUN (bottom) strains. Each row represents one sequence read and each column one cytosine. Blue squares represent converted cytosines, red squares represent unconverted cytosines, sequencing mismatches are shown in light blue and gaps are shown in white. 3 independent biological replicates. **(B)** Percent identity matrix of human NSUN proteins according to the Clustal Omega multiple alignment tool. Image generated using GraphPad Prism utilising values generated from alignment analysis. **(C, D)** Targeted bisulfite-sequencing heat map showing non-conversion rates of cytosines in mitochondrial tRNA Met-CAU (C) and mitochondrial 18S rRNA (D). Each row represents one genetic strain analysed and each column represents one cytosine. 2 independent biological replicates, 10 clones sequenced per strain, per replicate.

A BLAST analysis of the human NSUN3 methyltransferase domain against the *C. elegans* proteome showed higher similarity to NSUN-4 (29.93% identity), followed by NSUN-2 (26.13% identity) (data not shown). Moreover, we observed that, among human NSUN genes, NSUN3 and NSUN4 share the highest percentage of similarity (**Figure 3B**). Using a targeted BS-seq approach, we probed the methylation status of position C34 in mitochondrial tRNA Met-CAU from wild type, *nsun-2* and *nsun-4* strains. Our results indicate that NSUN-4 is responsible for the catalysis of m^5^C in this position in *C. elegans* (**Figure 3C**).

NSUN-4 is the only mitochondrial rRNA m^5^C methyltransferase identified to date. To confirm that the previously reported role of NSUN4 is also conserved in *C. elegans*, we performed targeted BS-seq in 18S mitochondrial rRNA, and found that methylation of positions C628 and C632, as well as C631, is mediated by NSUN-4 (**Figure 3D**). Notably, methylation of position C631 was also detected by WTBS, however in reduced stoichiometry (23.5% in wild type vs. 0.04% in noNSUN). Taken together, our results show that NSUN-4 is a multisite-specific tRNA/rRNA mitochondrial methyltransferase in nematodes.

### Loss of m^5^C leads to temperature-sensitive developmental and reproductive phenotypes

The individual or collective introduction of mutations in *nsun* genes failed to induce noticeable gross abnormal phenotypes. We therefore performed a more extensive characterisation of the mutant strains using a live imaging-based phenotypic analysis (Akay et al., 2019). As a proxy for reduced fitness, we chose to analyse the number of viable progeny and occurrence of developmental delay (growth rate, as measured by body length).

In comparison to wild type animals, we observed a reduction in body length in all mutant strains, which persists throughout development and into adulthood. This difference is greater in noNSUN animals, especially as this strain transitions from L4 stage to young adulthood (**Figure 4A**). In addition, the noNSUN strain showed a 25% reduction in brood size, which is comparable to what is observed in *nsun-1* and *nsun-5* individual mutants (**Figure 4B**).

**Figure 4.**
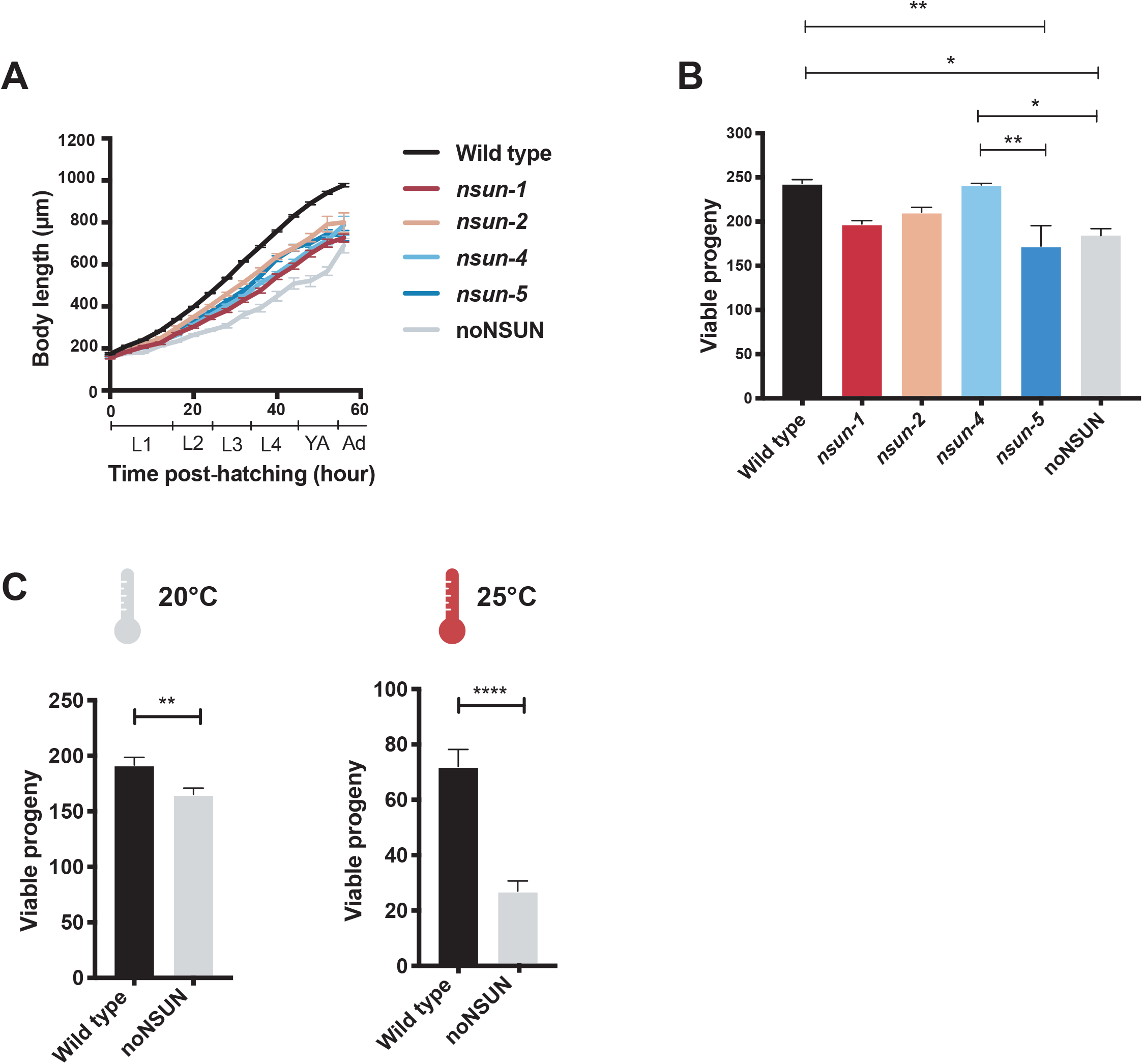
Loss of m^5^C leads to a temperature-sensitive reproductive phenotype. **(A)** Body length of individual *nsun* mutants throughout development shown as the median length of all individual animals (n=44,7,7,7,8,8) in ∼4 hour windows, error bars indicate the 95% confidence interval of the median, L1-L4 refers to the larval stages, YA and Ad to young adult and adult, respectively. (**B, C)** Viable progeny counts of individual *nsun* mutants at 20°C (B) and of noNSUN strain at different environmental temperatures (C). 10 young adult nematodes of each genotype were allowed to lay eggs for 4 days. Viable progeny was either manually or automatically counted at young adult stage. Automatic counting was done using using a Matlab script which processed plate images in real-time. Mean with SEM; One-way ANOVA, Tukey’s test.

*C. elegans* stocks can be well maintained between 16 °C and 25 °C, being most typically kept at 20 °C. To gain insights into how the loss of m^5^C impacts development under different environmental conditions, wild type and noNSUN animals were cultured at 25 °C for three generations and subjected to automated measurements. As shown in **Figure 4C**, the reproductive phenotype previously observed in the noNSUN strain (**Figure 4B**) is significantly aggravated at this temperature. More specifically, a reduction of 14% in brood size at 20 °C is worsened to 62.5% at 25 °C. This suggests that the phenotypes arising from loss of m^5^C are temperature-sensitive, pointing towards an involvement of this modification in the adaptation to environmental changes.

### Loss of m^5^C impacts decoding efficiency of leucine and proline codons

To explore the impact of temperature stress in the absence of m^5^C while avoiding the confounding effect introduced by differences in brood size, we performed further experiments using an acute heat shock treatment. To investigate whether the observed phenotypes are linked to abnormalities in protein translation rates, we quantified the polysomal fraction in wild type and noNSUN adult animals subject to heat shock at 27 °C for 4 hours, and found no significant differences (**Figure 5A**).

**Figure 5.**
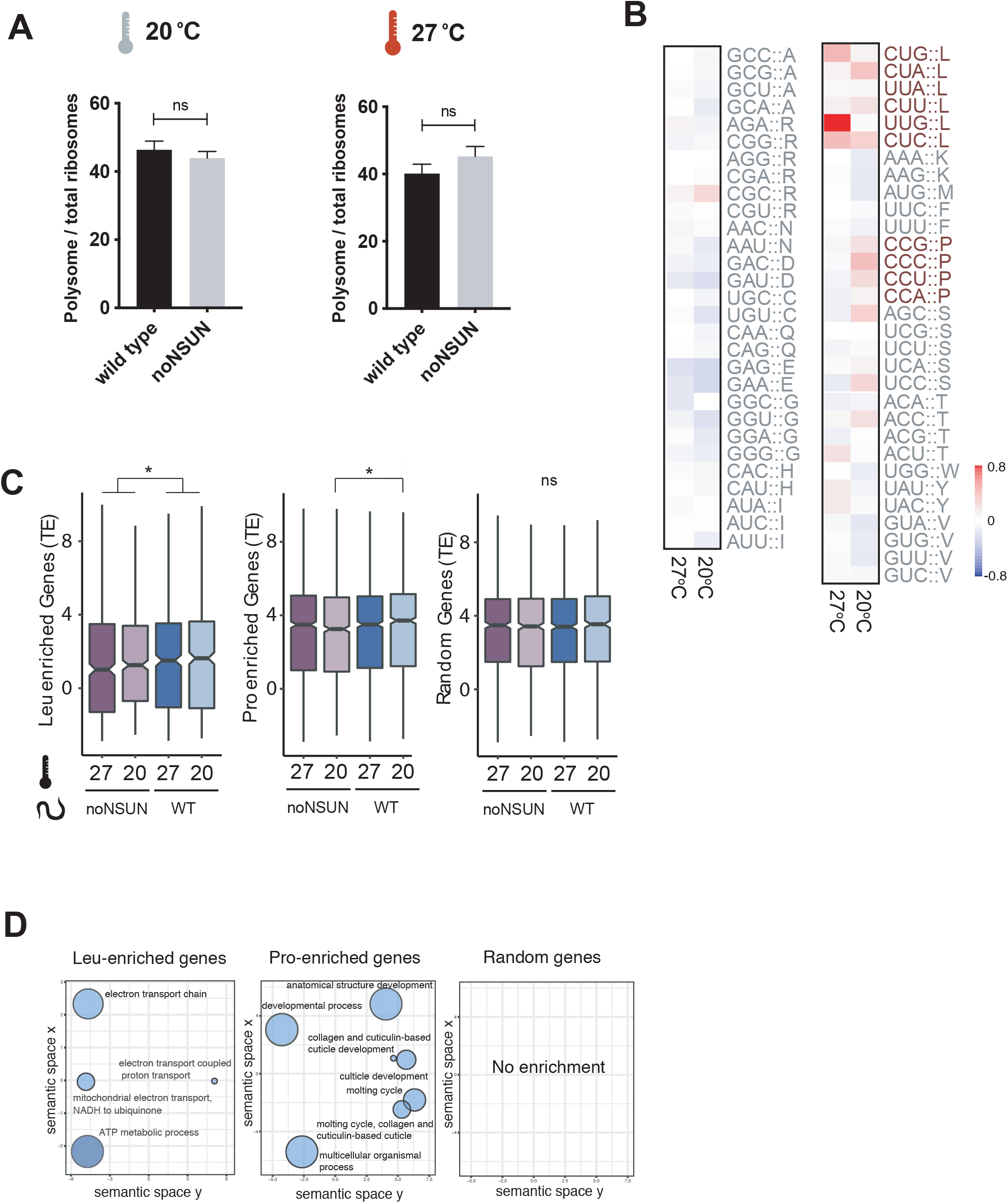
Loss of m^5^C impacts translation efficiency of leucine and proline codons. **(A)** Fraction of polysomal ribosomes quantified from polysome profiles from the wild type and noNSUN strains subject to a 4 h heat shock at 27°C. ns = non-significant. **(B)** Log2 fold change of bulk codon occupancy between wild type and noNSUN at 20°C and 27°C for each codon. (**C)** Translation efficiency of leucine-enriched, proline-enriched and random genes in each sample. (**D)** Gene ontology enrichment (biological process) of leucine-enriched, proline-enriched and random genes. 3 biological replicates.

To gain insights into transcriptional and translational differences resulting from the loss of m^5^C, we performed transcriptomic and ribosome profiling analyses. The latter allows the quantification of active translation by deep-sequencing of the mRNA fragments that are protected from nuclease digestion due to the presence of ribosomes (ribosome protected fragments - RPFs). RPFs showed the expected 3 nt periodicity along the coding domain sequences of mRNA, with the majority of reads in frame to the CDS start site (**Supplementary Figure 4A and 4B**). Furthermore, RNASeq and riboSeq counts of genes showed high correlation, and variation in the gene counts could be attributed to the difference in samples analysed (**Supplementary Figure 4C, 4D and 4E**). Loss of m^5^C did not greatly impact the nature of the heat stress response, as most differentially transcribed and translated genes upon heat stimulus showed agreement, or very subtle differences between wild type and noNSUN strains (**Supplementary Figure 5**). We found that differentially transcribed genes upon loss of m^5^C are mainly involved in cuticle development, while differentially translated genes are enriched in components of the cuticle and ribosomes, as well as RNA-binding proteins (**Supplementary Figure 6**).

We then evaluated genome-wide codon occupancy during translation elongation and found that loss of m^5^C leads to increased ribosome occupancy at leucine and proline codons. More specifically, upon heat shock Leu-UUG codons showed the highest ribosome density observed in the noNSUN strain, suggesting that translation of this codon is slowed upon heat stress in the absence of m^5^C (**Figure 5B**). Interestingly, as shown in our WTBS analysis, leucine and proline are the most frequently methylated tRNA isoacceptors in *C. elegans* (**Figure 2B**). In addition, tRNA Leu-CAA, responsible for decoding of UUG codons, is the only cytoplasmic tRNA bearing an m^5^C-modified wobble position (**Figure 2C**). Corroborating these findings, we found translation efficiency of leucine and proline-enriched genes to be significantly reduced in the noNSUN strain. This effect is more pronounced in leucine-rich genes and is observed in both temperatures tested, becoming more pronounced as transcripts get more enriched in this amino acid (**Figure 5C**, **Supplementary Figure 7**). Finally, we found that translationally affected leucine-enriched genes are mainly involved in mitochondrial functions, while proline-enriched genes are involved in anatomical development (**Figure 5D**).

## DISCUSSION

Chemical modifications of RNA occur in organisms from all kingdoms of life and are often highly conserved throughout evolution, as is the case of the methylation of carbon-5 in cytosines (Boccaletto et al., 2017). While this is the first report of an animal completely devoid of m^5^C, our results add to the growing body of evidence showing that several RNA modifications are individually not required for development under controlled conditions ((O’Connor, Leppik, & Remme, 2018), reviewed in (Sharma & Lafontaine, 2015) and (Hopper & Phizicky, 2003)). *In vitro*-transcribed rRNA and tRNA, *i.e.* non-modified, can be successfully used for ribosomal assembly and reconstituted translation assays (Cload, Liu, Froland, & Schultz, 1996; Green & Noller, 1999; Khaitovich, Tenson, Kloss, & Mankin, 1999; Semrad & Green, 2002), suggesting that modifications are often not essential for the basic roles executed by these non-coding RNAs in translation. Our results reignite a recurrent question in the epitranscriptomics field: why are so many of these chemical marks extensively conserved throughout evolution and, yet, organisms seem to develop normally in their absence? Ribonucleoside modifications occur in an overwhelming diversity and, in some cases, are likely to *(i)* exert subtle molecular effects, *(ii)* act in a combinatorial manner with other modifications or *(iii)* be the result of relaxed enzymatic specificity (Jackman & Alfonzo, 2013; Phizicky & Alfonzo, 2010).

Our results suggest an RNA methylation-independent essential role for NSUN-1 in germline development. Overall, Nop2p/Nol1/NSUN1 has been shown to be an essential gene also in yeast, mice and *Arabidopsis* (Burgess et al., 2015; De Beus, Brockenbrough, Hong, & Aris, 1994; Kosi et al., 2015; Sharma et al., 2013). In *Saccharomyces cerevisiae*, temperature-sensitive alleles, depletion of Nop2p, and mutations in catalytic residues, all cause defective rRNA processing and lower levels of 60S ribosomal subunits, supporting the idea that these molecular phenotypes are linked to reduced methylation (Hong, Brockenbrough, Wu, & Aris, 1997; Hong, Wu, Brockenbrough, Wu, & Aris, 2001; Sharma et al., 2013). In contrast, Bourgeois et al (Bourgeois et al., 2015) reported that loss of m^5^C in position 2870 of helix 89 of 25S rRNA had no effect on ribosome synthesis, nor does it cause a phenotype. Therefore, the role NSUN-1-mediated m^5^C sites remain to be determined. A similar phenomenon has been observed in NSUN4 mutant mice and Dim1 mutant yeast, where the presence of the enzyme, rather than its catalytic activity, is required for viability (26,63,64). It has been suggested that the essential binding of certain rRNA methyltransferases represents a quality control step in ribosome biogenesis, committing rRNA to methylation during the maturation process (D. L. Lafontaine, Preiss, & Tollervey, 1998).

Taking advantage of the noNSUN strain as a tool to increase the confidence of WTBS analysis, we produced the first comprehensive list of m^5^C sites throughout *C. elegans* transcriptome. Using a targeted approach, we showed that NSUN-4 has both rRNA and tRNA targeting capabilities in the mitochondria. It has been suggested that NSUN4 lacks an RNA recognition domain and would, therefore, need a binding partner in order to achieve targeting (Cámara et al., 2011). Crystal structure analyses showed that the MTERF4-NSUN4 complex forms a positively charged path along the sides of MTERF4, which extends into the active site of NSUN4. This structure was found likely to be responsible for targeting this methyltransferase to rRNA in the mitochondria, thus providing the sequence specificity for RNA binding (Spåhr et al., 2012; Yakubovskaya et al., 2012). Nevertheless, genetic evidence suggests that NSUN4 methylates rRNA independently of MTERF4 in mice (Metodiev et al., 2014). *C. elegans* has a MTERF4 homologue (K11D2.5), and most residues involved in the interaction between NSUN4 and MTERF4 are conserved or show convergent evolution, suggesting that this interaction could occur in this organism (Spåhr et al., 2012). More experiments will be required to clarify the mechanisms through which NSUN-4 shows multisite targeting capabilities in the nematode.

In humans, NSUN3-mediated methylation at position 34 of mitochondrial tRNA Met-CAU is further modified by the dioxygenase ALKBH1 to form f^5^C (Haag et al., 2016; Nakano et al., 2016). Additional analysis, such as specific isolation of tRNA Met-CAU using DNA probes, have been previously used in *C. elegans* RNA and successfully confirmed the presence of f^5^C in this organism (Nakano et al., 2016). As f^5^C reacts as an unmodified cytosine upon sodium bisulfite treatment, it was surprising to detect such high levels of non-conversion (95%) in the present study. Previous studies tackling this question explored differential methods for the detection of f^5^C, such as reduced bisulfite sequencing (RedBS-Seq), f^5^C chemically assisted bisulfite sequencing (fCAB-Seq) and LC/MS of isolated mt-tRNA Met-CAU. Their results indicate that 35-100% of tRNA Met-CAU molecules are f^5^C-modified, while the whole population is at least m^5^C-modified (Haag et al., 2016; Kawarada et al., 2017; Van Haute et al., 2016). As we used a sequencing-based method that does not select for precursors or mature tRNAs, it is possible that our results are biased towards the detection of primary transcripts or precursor molecules, which may not have been oxidised by ALKBH1 yet.

Using the noNSUN strain as a negative control and as a proxy for methylation-derived non-conversion of cytosines, we investigated the presence of m^5^C in coding transcripts. Several reports have supported the idea that m^5^C is a common mRNA modification (Amort et al., 2017; David et al., 2017; Squires et al., 2012; Yang et al., 2017). Results derived from BS-seq can be influenced by several sources of artefacts, such as incomplete deamination, protection due to secondary structures, presence of other modifications, protein binding, sequencing errors, among others (summarized in (Legrand et al., 2017)). Given these technical drawbacks, the noNSUN strain represented an unprecedentedly stringent negative control, which allowed for exclusive detection of highly specific methylation. Despite the detection of several positions with 20-30% NSUN-dependent non-conversion, we detected a similar number of positions with NSUN-independent non-conversion at these rates, which we interpret as false positives. This poses a statistical challenge on the interpretation of such non-converted positions as methylated. Our main conclusion, therefore, is that the data does not provide evidence for widespread or high stoichiometry m^5^C methylation of coding transcripts in *C. elegans*. This is in agreement with earlier work using chromatography (Adams & Cory, 1975; Desrosiers, Ronald Friderici, Karen Rottman, 1974; Salditt-Georgieff et al., 1976) and other reports that have detected very few or no m^5^C methylated sites in eukaryotes by BS-seq (Edelheit, Schwartz, Mumbach, Wurtzel, & Sorek, 2013; Legrand et al., 2017).

While the absence of m^5^C in RNA did not give rise to overt phenotypes under standard laboratory conditions, a more detailed analysis of the mutants revealed developmental and fertility defects. In nature, where the environmental conditions vary greatly, such genotypes might be selected against in a wild population. In fact, previous studies have shown that levels of several RNA modifications, including m^5^C, are responsive and can react dynamically to a wide range of environmental challenges, such as toxicants, starvation and heat shock, thus potentially supporting organismal adaptation (C. T. Y. Chan et al., 2010; van Delft et al., 2017). In agreement with this idea, we observed a temperature-dependent aggravation of reproductive phenotypes in m^5^C-deficient *C. elegans*.

Despite the fact that our ribosome profiling analysis was performed in a mutant strain lacking m^5^C in all types of RNAs, the effect observed on translational speed was strikingly specific. The strongest effect was observed in UUG codons, which are dependent on the only tRNA modified at the wobble position, tRNA Leu-CAA. Chan et al. (C. T. Y. Chan et al., 2012) found that m^5^C level specifically at position 34 of tRNA Leu-CAA is upregulated upon oxidative stress in yeast. The presence of this modification was shown to enhance translation efficiency of a UUG-rich luciferase reporter construct, as *trm4*Δ (NSUN2 homologue mutant) cells showed significantly lower levels of reporter activity, especially under oxidative stress. The biological relevance of these findings was linked to an abnormally high frequency of UUG codons in transcripts of specific ribosomal protein paralogues. Our results suggest that the genes potentially affected by slowed translation of leucine codons are involved in the electron transport chain (**Figure 5C**). Interestingly, a recent report demonstrated that NSUN2 mutant cells show differential expression of mitochondrial genes, and contain less-active mitochondria upon stress induced by sodium arsenite treatment (Gkatza et al., 2019).

In summary, our work reveals the biology and mechanism of m^5^C deposition in *C. elegans*, providing a list of high-confidence m^5^C sites in the nematode’s transcriptome, as well as their enzymatic specificity. While this RNA modification is not essential for viability under laboratory conditions, its presence is associated with increased fitness at a higher temperature, and enhanced translational efficiency of leucine and proline codons. The most pronounced effect was observed for UUG codons exclusively upon heat shock, suggesting a role of m^5^C tRNA wobble methylation in the adaptation to heat stress (**Figure 6**).

**Figure 6.**
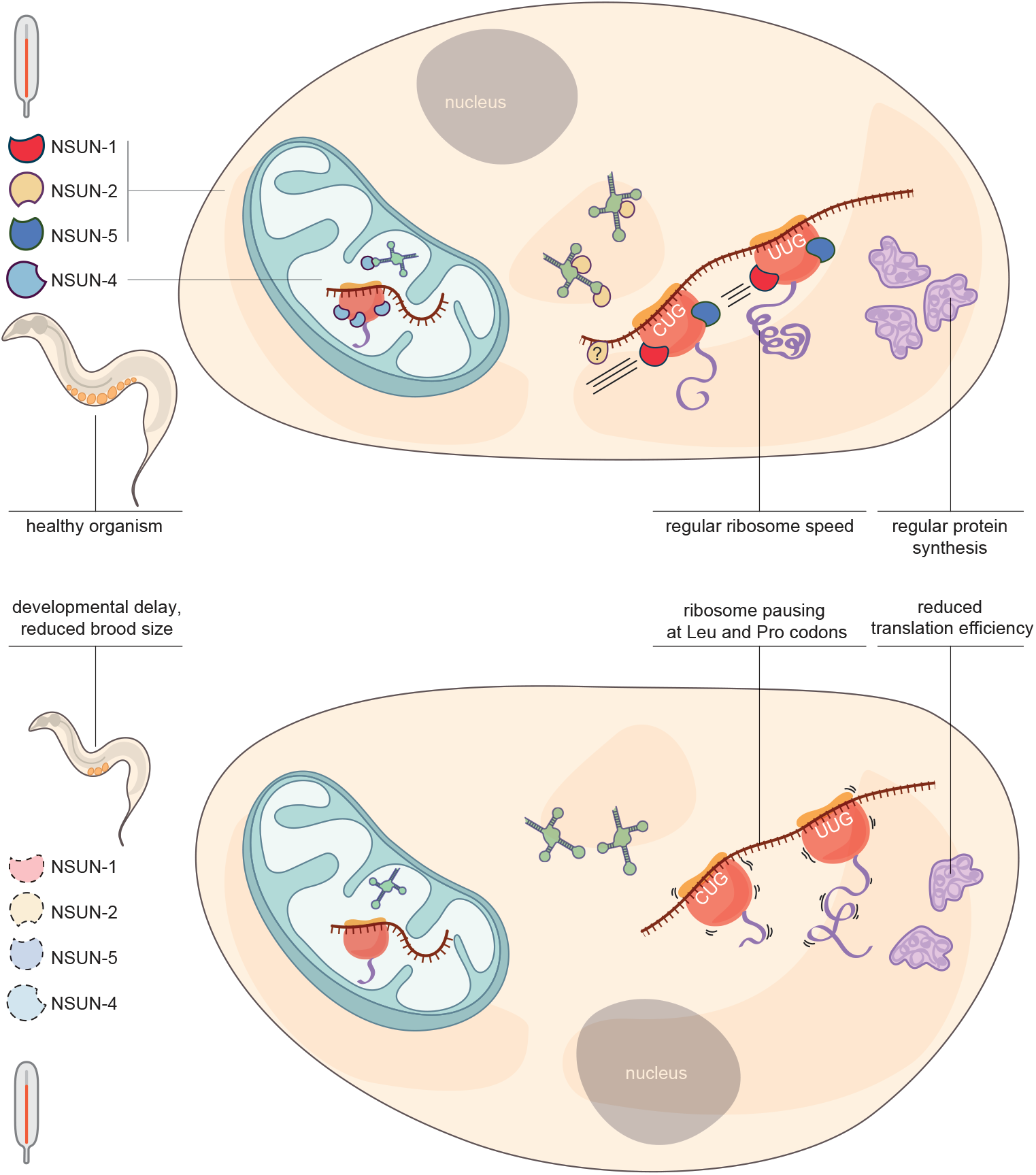
The m^5^C pathway of *C. elegans*. Top: The nematode’s genome encodes four m^5^C-methyltransferases: NSUN-1, NSUN-2, NSUN-4 and NSUN-5 (here represented in red, yellow, light blue and dark blue, respectively). In the cytoplasm, NSUN-1 and NSUN-5 modify one position in rRNA each, while NSUN-2 modifies the variable loop of several tRNAs and the wobble position of tRNA Leu-CAA. In the mitochondria, NSUN-4 modifies 3 positions in rRNA and the wobble position of tRNA Met-CAU. In the wild-type strain, ribosomes move at regular speed to achieve regular translation efficiency. Bottom: Simultaneous mutation of all m^5^C-methyltransferases completely abolishes m^5^C presence in RNA and leads to developmental delay and reduced brood size, which is aggravated at higher temperature. In the noNSUN strain, ribosomes tend to pause at the encounter with leucine and proline codons, leading to reduced translation efficiency of transcripts enriched in these amino acids.

## MATERIALS AND METHODS

### Genetics

*C. elegans* strains were grown and maintained as described in Brenner (Brenner, 1974). The strains were kept at 20 °C, unless otherwise indicated. HB101 strain *Escherichia coli* was used as food source (*Caenorhabditis* Genetics Center, University of Minnesota, Twin Cities, MN, USA). Bristol N2 was used as the wild type strain.

### Gene silencing by RNAi

Empty vector, *nsun-1* (*W07E6.1*), *nsun-2* (*Y48G8AL.5*), *nsun-4* (*Y39G10AR.21*), and *nsun-5* (*Y53F4B.4*) bacterial feeding clones were kindly provided by Prof. Julie Ahringer’s lab (Kamath et al., 2003). Single colonies were inoculated in LB-Ampicillin 100 μg/ml and cultured for 8 h at 37 °C. Bacterial cultures were seeded onto 50 mm NGM agar plates containing 1 mM IPTG and 25 μg/ml Carbenicillin at a volume of 200 μl of bacterial culture per plate, and left to dry for 48 hours. 50 synchronized L1 larvae were placed onto RNAi plates and left to grow until adult stage. Adults were scored for fertility (presence of embryos in the germline).

### CRISPR-Cas9 gene editing

CRISPR-Cas9 gene editing was performed as in Paix et al (Paix, Folkmann, Rasoloson, & Seydoux, 2015). Briefly, injection mixes were prepared in 5 mM Tris pH 7.5 as follows: 20 μg of tracrRNA (Dharmacon), 3.2 μg of *dpy-10* crRNA (Dharmacon), 200 ng of *dpy-10* homologous recombination template (Sigma Aldrich), 8 μg of target gene gRNA (Dharmacon), 1.65 μg of homologous recombination template (Sigma), up to a volume of 11.5 μl. The mix was added to 10 μg of Cas9 (Dharmacon) to a final volume of 15 μl, and incubated at 37 °C for 15 min. For the creation of *nsun* catalytic mutants, a homologous recombination template bearing a point mutation to convert the catalytic cysteine into alanine while creating a restriction site for HaeIII was co-injected. Following incubation, the mix was immediately micro-injected into the germline of N2 young adults. After injection, animals were left to recover in M9 medium, then transferred to individual plates and left to recover overnight at 20 °C. Successful injections led to the hatch of dumpy and roller animals. From positive plates, 96 animals were individualized for self-fertilization and genotyped for the relevant alleles. Same process was performed with F2s, until a homozygous population was isolated. Each strain was backcrossed at least three times with the wild type strain.

### RNA extraction (Mass spectrometry and WTBS)

The strains of interest (N2 and noNSUN) were grown in 90 mm plates until gravid adult stage, washed three times with M9 and pelleted by centrifugation at 2000 rpm for 2 minutes. Gravid adults were resuspended in 4 ml of bleaching solution (final concentration 177 mM NaOH, 177 mM NaOCl solution - free chlorine 4–4.99%) and vortexed vigorously for 7 minutes. Recovered embryos were washed four times to remove any traces of bleach and left to hatch in ml of M9 for 24 h at 20 °C in a rotating wheel. Synchronised L1 starved larvae were used for RNA extraction. Independent triplicates were obtained from three different generations. Nematodes were washed thoroughly in M9 to remove bacterial residue, and pelleted in RNAse-free tubes at 2000 rpm for 2 min. 500-1000 μl of TRIsure (Bioline) and 100 μl of zirconia beads were added and the samples were subjected to three cycles of 6,500 rpm with 20 sec breaks on Precellys to crack open the animals. 100 μl of chloroform were added to the tubes, which were then shaken vigorously for 15 sec and incubated at room temperature for 3 min. Samples were centrifuged at 12,000 x *g* for 15 min at 4 °C and the aqueous phase of the mixture was carefully recovered and transferred to a fresh RNAse-free tube. RNA was precipitated with 500 μl of cold isopropanol at room temperature for 10 min and then centrifuged at 12,000 x *g* for 15 min at 4 °C. The supernatant was carefully removed, the pellet was washed and vortexed with 1 ml of 75% ethanol and centrifuged at 7,500 x *g* for 5 min at 4 °C. RNA pellet was air-dried, dissolved in the appropriate volume of DEPC-treated water and the concentration, 260/280 and 260/230 ratios were measured by Nanodrop. RNA integrity was evaluated in the Agilent 2200 Tapestation system.

### RNA mass spectrometry

Up to 10 μg of RNA was digested by adding 1 μl digestion enzyme mix per well in a digestion buffer (4 mM Tris-HCl pH 8, 5 mM MgCl2, 20 mM NaCl) in a total volume of up to 100 μl. The digestion enzyme mix was made by mixing benzonase (250 U/μl, Sigma Aldrich), phosphodiesterase I from *Crotalus adamanteus* venom (10mU/μl, Sigma Aldrich) and Antarctic phosphatase (5 U/μl, NEB) in a ratio of 1:10:20. The reaction was incubated overnight at 37 °C. The following day, an equal volume of ^13^C, ^15^N-labelled uridine (internal control, previously dephosphorylated; Sigma Aldrich) in 0.1% formic acid was added to each reaction and this was subsequently prepared for LC-MS-MS by filtration through 30 kDa molecular weight cut-o filters (Sigma).

Samples were resolved using a Thermo Scientific U3000 UPLC system on a gradient of 2-98% (0.1% formic acid/acetonitrile) through an Acquity 100mm x 2.1 mm C-18 HSS T3 column and analysed on a QExactive-HF Orbitrap High Resolution Mass Spectrometer (ThermoFisher Scientific, IQLAAEGAAPFALGMBFZ) in positive full-scan mode and the results were deconvoluted using the accompanying Xcalibur Software. Nucleosides of interest were identified by both retention times and accurate masses, compared to purified standards and quantified accordingly.

### Whole Transcriptome Bisulfite Sequencing

Bisulfite sequencing experiments were performed as previously described in Legrand et al. (Legrand et al., 2017). RNA was fractionated into <200 nt and >200 nt using a modified mirVana miRNA isolation kit (AM1560) protocol. Briefly, 50 μg of RNA in a volume of 80 μl were mixed with 400 μl of mirVana lysis/binding buffer and 48 μl of mirVana homogenate buffer and incubated for 5 min at room temperature. Next, 1/3 volume (176 μl) of 100% ethanol was added and thoroughly mixed by inversion, and the mixture was incubated for 20 min at room temperature. After addition of 0.8 μg of Glycoblue, the samples were spun down at 2,500 x g for 8 min at 21 °C for precipitation of long RNAs. The supernatant containing the short fraction was transferred to a fresh tube and the RNA pellet was washed in 1 ml of cold 75% ethanol before centrifugation at maximum speed for at least 20 min at 4 °C. The pellet was finally air-dried and resuspended in DEPC-treated water. For short fraction RNA precipitation, 800 μl of isopropanol were added to the supernatant and the mixture was incubated at −80 °C for at least 20 min. Next, 20 μg of Glycoblue were added and the mixture was spun down at maximum speed for at least 20 min at 4 °C. The pellet was washed with cold 70% ethanol and air dried before resuspension in DEPC-treated water. Depletion of ribosomal RNA was performed on the short fractions and on half of the long fractions using a Ribo-zero rRNA removal kit (Illumina), according to the supplier’s instructions. The other half of long fractions was processed as Ribo+ samples. RNA was stored at −80 °C until the moment of use.

The long fractions (with and without rRNA depletion) were further processed with the NEBNext Magnesium RNA Fragmentation Module (NEB), as described in the manual. 3 min of fragmentation at 94 °C has been established to lead to a peak at approximately 250 nt, appropriate for the final 100 bp paired-end sequencing. The fragmented RNA was precipitated using ethanol with 20 μg GlycoBlue at −80 °C for at least 10 min.

Samples were treated with TURBO DNase (Ambion) in a final volume of 20 μl, according to the manufacturer’s instructions. DNase-treated samples were bisulfite-converted using an EZ RNA Methylation Kit (Zymo Research), following the manufacturer’s manual. As a final step before library preparation, a stepwise RNA end repair was carried out using T4 polynucleotide kinase (TaKaRa). A 3′-dephosphorylation and 5′-phosphorylation reaction was performed using T4 PNK enzyme (TaKaRa). The enzyme was removed by phenol-chloroform purification. Library preparation was done using a NEBNext Small RNA Library Prep Set, according to the manufacturer’s protocol. cDNA was amplified with 12 cycles of PCR and purified using the QIAquick PCR Purification Kit (Qiagen). The libraries were size-selected on a 6% polyacrylamide gel. Compatible barcodes were selected, and samples were pooled in equimolar ratios on multiple lanes in an Illumina HiSeq 2000 platform. A 100 bp paired-end sequencing approach was used.

Bioinformatics, statistical analyses and methylation calling were performed as described in Legrand et al. (Legrand et al., 2017), utilising the BisRNA software. Adapters were removed from sequenced reads using Cutadapt version 1.8.1 (with options: --error-rate=0.1 --times=2 --overlap=1 and adapter sequences AGATCGGAAGAGCACACGTCT and GATCGTCGGACTGTAGAACTCTGAAC for forward and reverse reads, respectively (Martin, 2011). Reads were further trimmed of bases with phred quality score <30 on 5’ and 3’ ends and reads shorter than 25 nucleotides were discarded (Trimmomatic version 0.36) (Bolger, Lohse, & Usadel, 2014). Reads were aligned uniquely using Bsmap (version 2.87, options: -s 12 -v 0.03 -g 0 -w 1000 -S 0 -p 1 -V 1 -I 1 -n 0 -r 2 -u -m 15 -x 1000) (Xi & Li, 2009). Reference sequences were downloaded from Gtrnadb (version ce10), Ensembl (release 90, version WBcel235) and Arb-Silva (P. P. Chan & Lowe, 2016; Kersey et al., 2016; Lee et al., 2018; Quast et al., 2012). End sequence ‘CCA’ was appended to tRNA if missing. Bisulfite-identical sequences, where only C>T point differences were present, were merged, keeping the C polymorphism. Similarly to Legrand et al. (Legrand et al., 2017), tRNA sequences were further summarized to the most exhaustive yet unambiguous set of sequences, using sequence similarity matrix from Clustal Omega (Sievers et al., 2011). Methylation calling were performed as described in Legrand et al. (Legrand et al., 2017), utilising the BisRNA software. Methylation frequency was calculated as the proportion of cytosines with coverage higher than 10 in three wild type and noNSUN replicates and bisulfite non-conversion ratio higher than 0.1. The measure for reproducibility was the standard error. Deamination rates were calculated as the count of converted cytosines divided by the sum of converted and non-converted cytosines. This calculation was carried out on nuclear and mitochondrial rRNA. Known methylation sites in rRNA were removed from the calculations. WTBS raw data have been deposited in the Gene Expression Omnibus (GEO) database under the accession number GSE144822.

### Targeted bisulfite sequencing

1 μg of total RNA was bisulfite-modified with the EZ RNA Methylation Kit (Zymo Research). Briefly, samples were first treated with DNase I for 30 min at 37 °C in 20 μl volume. The DNAse reaction was stopped and immediately applied to the EZ RNA Methylation Kit (Zymo Research) according to manufacturer’s instructions. Converted RNAs were eluted in 12 μl of distilled water.

Reverse transcription was performed with the purified RNA and adaptors were added to the amplicons using reverse oligonucleotides designed for the bisulfite-converted sequences of interest (reverse adaptor sequence: gtctcgtgggctcggagatgtgtataagagacag) and SuperScript III reverse transcriptase (Invitrogen). cDNA was cleaned from any residual RNA with an RNase H treatment at 37 °C for 20 min and then used for PCR amplification and adaptor addition using forward oligonucleotides (forward adaptor sequence: tcgtcggcagcgtcagatgtgtataagagacag). Low annealing temperature (58 °C) was used to overcome high A-T content after bisulfite treatment. 4 μl of PCR product were used for ligation and transformation into TOP10 competent cells using the Zero Blunt TOPO PCR Cloning Kit (Invitrogen) according to the manufacturer’s instructions. Following overnight culture, 24 colonies were individually lysed and used for PCR amplification using M13 primers, in order to confirm the presence of the insert at the correct size by DNA electrophoresis. The remaining PCR product (10 clones per condition) was used for Sanger sequencing using T3 primers (Genewiz).

### Automated phenotypical characterisation

#### Viable Progeny

Viable progeny refers to the number of progeny able to reach at least the L4 stage within ∼4 days. Measurements were completed over three 24-h intervals. First, eggs were prepared by synchronisation via coordinated egg-laying. When these animals had grown to the L4 stage, single animals were transferred to fresh plate (day 0). For 3 days, each day (days 1–3), each animal was transferred to a new plate, while the eggs were left on the old plate and allowed to hatch and grow for ∼ 3 days, after which, the number of animals on each of these plates was counted (Hodgkin & Barnes, 1991) using a custom animal counting program utilising short video recordings. Animals were agitated by tapping each plate four times, after this, 15 frames were imaged at 1 Hz and the maximum projection was used as a background image. Animals were then detected by movement using the difference in the image between each frame and this background image and counted this way for ten additional frames. The final count was returned as the mode of these counts. This system was tested on plates with fixed numbers of animals and was accurate to within 5%, comparable to human precision. Total viable progeny was reported then as the sum for 3 days. Data is censored for animals that crawled off of plates.

#### Single Worm Growth Curves

Populations of *Caenorhabditis elegans* were synchronized by coordinated egg laying. Single eggs were transferred to individual wells of a multi-well NGM plate solidified with Gelrite (Sigma). Each well was inoculated with 1 *μl* of OD 20 *E. coli* HB101 bacteria (∼18 million) and imaged periodically using a camera mounted to a computer controlled XY plotter (EleksMaker, Jiangsu, China) which moved the camera between different wells. Images were captured every ∼11 minutes for ∼75 hours. Image processing was done in real-time using custom MATLAB scripts, storing both properties of objects identified as *C. elegans*, and sub images of regions around detected objects. Body length was calculated using a custom MATLAB (Mathworks, Natick MA) algorithm and all other properties were measured using the *regionprops* function. Growth curves were aligned to egg-hatching time, which was manually determined for each animal.

### Polysome profiling

Synchronised populations of the strains of interest (N2 and noNSUN) were grown until adult stage (3 days) in 140 mm NGM agar plates seeded with concentrated *E. coli* HB101 cultures at 20 °C. Next, the animals were harvested from the plates, transferred to liquid cultures in S-medium supplemented with *E. coli* HB101, and incubated at 20°C or 27°C for 4h in a shaking incubator at 200 rpm before harvesting. Sample preparation for polysome profiling was adapted from Arnold et al (Arnold et al., 2014). The animals were harvested, washed 3x in cold M9 buffer supplemented with 1 mM cycloheximide and once in lysis buffer (20 mM Tris pH 8.5, 140 mM KCl, 1.5 mM MgCl2, 0.5% Nonidet P40, 2% PTE (polyoxyethylene-10-tridecylether), 1% DOC (sodiumdeoxycholate monohydrate), 1 mM DTT, 1 mM cycloheximide). The animals were pelleted and as much liquid as possible was removed before the samples were frozen as droplets in liquid nitrogen, using a Pasteur pipette. Frozen droplets were transferred to metallic capsules and cryogenically ground for 25 sec in a mixer (Retsch MM 400 Mixer Mill). The resulting frozen powder was stored at −80 °C until the moment of use.

Approximately 250 μl of frozen powder was added to 600 μl of lysis buffer and mixed by gentle rotation for 5 min at 4 °C. The samples were centrifuged at 10,000 x g for 7.5 min, the supernatant was transferred to fresh tubes and the RNA concentration was quantified by Nanodrop. For ribosome footprinting, 400 μl of lysate was treated with 4 μl of DNase I (1 U/μl, Thermo Scientific) and 8 μl of RNase I (100 U/μl, Ambion) for 45 min at room temperature with gentle shaking. 20 μl of RNasin ribonuclease inhibitor (40 U/μl, Promega) was added to quench the reaction when appropriate. The tubes were immediately put on ice and 220 μl of lysate was loaded into 17.5 – 50% sucrose gradients and ultracentrifuged for 2.5 h at 35,000 rpm, 4 °C in a Beckman SW60 rotor. In parallel, undigested samples used for polysome profiling were equally loaded into sucrose gradients under the same conditions. Gradient fractions were eluted with an ISCO UA-6 gradient fractionator while the absorbance at 254 nm was continuously monitored. The fraction of polysomes engaged in translation was calculated as the area under the polysomal part of the curve divided by the area below the entire curve.

### Ribosome profiling

Sucrose gradient fractions were collected in tubes containing 300 μl of 1 M Tris-HCl pH 7.5, 5M NaCl, 0.5 M EDTA, 10% SDS, 42% urea and then mixed by vortexing with 300 μl of Phenol-Chloroform-Isoamylalcohol (PCL, 24:25:1). Fractions corresponding to 80S monosomes were heated for 10 min at 65°C and centrifuged at 16,000 x g for 20 min at room temperature. The upper aqueous phase was transferred to a fresh tube and mixed well with 600 μl of isopropanol and 1 μl of Glycoblue for precipitation overnight at −80 °C. RNA was pelleted by centrifugation at 16,000 x g for 20 min at 4 °C and washed in 800 μl of cold ethanol. After supernatant removal, the pellet was left to dry for 1-2 min and then dissolved in 60 μl of RNase-free water. RNA concentration and quality were measured by Nanodrop and Tapestation (2200 Agilent R6K), respectively.

A dephosphorylation reaction was performed by adding 7.5 μl T4 polynucleotide kinase (PNK) 10x buffer, 1.5 μl ATP, 1.5 μl RNase OUT, 1.5 μl T4 PNK (TaKaRa) and 3 μl RNase-free water up to a final volume of 75 μl, and incubated for 1.5 h at 37 °C. The enzyme was removed by acid-phenol extraction and the RNA was precipitated with 1/10 volume 3 M sodium acetate pH 5.2, 2.5x volume 100% cold ethanol and 1 μl Glycoblue overnight at −80 °C. Pelleted RNA was dissolved in RNase-free water. For footprint fragments purification, RNA was denaturated for 3 min at 70 °C and loaded into a 15% polyacrylamide TBE-urea gel alongside a small RNA marker. Gel was run for 1 h at 150 V and stained for 10 min with 1:10,000 SYBR Gold in 0.5x TBE. The gel was visualised under UV light and the region between the 20 nt and 30 nt marks (28-32 nt) was excised with a sterile scalpel. The gel band was crushed into small pieces and incubated in 300 μl of 0.3 M RNase-free NaCl solution with 2 μl of RNase OUT overnight at 4°C on an Intelli-Mixer (Elmi). The gel slurry was transferred to a 0.45 μm NanoSep MF Tube (Pall Lifesciences) and centrifuged at maximum speed for 5 min at 4°C. After overnight precipitation with 30 μl of 3 M sodium acetate pH 5.2, 1 μl Glycoblue and 800 μl 100% ethanol, the RNA was dissolved in RNase-free water.

Libraries were prepared using a NEB NEXT Small RNA Library Prep Set for Illumina (Multiplex compatible) E7330 Kit, following the manufacturer’s instructions. cDNA libraries were purified according to the manual, followed by a QIAQuick PCR Purification Kit and a 6% polyacrylamide gel, where a band of 150 bp (120 bp adapter + 28-32 footprint fragments) was excised. Gel extraction was performed as described above for footprint fragments purification. Libraries were sequenced at the Genomics and Proteomics Core Facility of the German Cancer Research Centre (DKFZ), Heidelberg.

Raw reads were assessed for quality using FastQC and Trimmed for low quality bases and adapter sequences using Trimmomatic (version 0.39, parameters - ILLUMINACLIP:2:30:10 SLIDINGWINDOW:4:20 MINLEN:20) (Bolger et al., 2014). SortMERNA was used to remove any rRNA sequences (Kopylova, Noé, & Touzet, 2012). Remaining reads were uniquely aligned to the *C. elegans* (WBCel235) reference genome using HISAT2 (version 2.1.0) (Kim, Langmead, & Salzberg, 2015). The longest transcript was chosen for each gene from the WBCel235 reference genome and the CDS for these transcripts was extracted. Reads were length stratified and checked for periodicity, only read lengths showing periodicity over the 3 frames were retained for further analysis (26 bp -30 bp). Reads aligned to the genome were shifted 12 bp from the 5′-end towards the 3′-end (Ingolia, Ghaemmaghami, Newman, & Weissman, 2009). Any reads aligned to the first 10 codons of each gene were then removed and the remaining reads with a 5′ end aligning to a CDS were kept for further analysis (Lecanda et al., 2016).

Bulk codon occupancy in the P-Site for each codon was calculated as the number of shifted RPFs assigned to the first nucleotide of the codon. This value was then normalized by the frequency of the counts for the same codon in the +1, +2 and +3 codons relative to the A-Site (Stadler & Fire, 2011). Fold changes were then computed as the normalized bulk codon occupancies for noNSUN / wild type. Ribosome occupancy for gene in a sample was calculated as the number of shifted in frame RPFs aligned to the CDS of the gene (not including the first 10 codons). These values were inputted into DESeq2 (Love, Huber, & Anders, 2014). Ribo-Seq raw data have been deposited in the Gene Expression Omnibus (GEO) database under the accession number GSE146256.

### RNA sequencing

Input RNA was extracted from aliquots from the samples used for polysome profiling and ribosome footprinting. 100 μl of chloroform were added to the tubes, which were then shaken vigorously for 15 sec and incubated at room temperature for 3 min. Samples were centrifuged at 12,000 x g for 15 min at 4 °C and the aqueous phase of the mixture was carefully recovered and transferred to a fresh RNAse-free tube. RNA was precipitated with 500 μl of cold isopropanol at room temperature for 10 min and then centrifuged at 12,000 x g for 15 min at 4 °C. The supernatant was carefully removed, the pellet was washed and vortexed with 1 ml of 75% ethanol and centrifuged at 7,500 x g for 5 min at 4°C. RNA pellet was air-dried, dissolved in the appropriate volume of DEPC-treated water and the concentration, 260/280 and 260/230 ratios were measured by Nanodrop. RNA integrity was evaluated in the Agilent 2200 Tapestation system. RNA was depleted of DNA with a TURBO DNA-free kit (Invitrogen), according to the manufacturer’s instructions. Libraries were prepared with 750 ng of starting material using the NEBNext Ultra II Directional RNA Library Prep Kit for Illumina, following rRNA depletion using a NEBNext rRNA Depletion Kit (Human/Mouse/Rat) (NEB).

Raw reads were assessed for quality using FastQC (**Andrews, 2010**) and Trimmed for low quality bases and adapter sequences using Trimmomatic (version 0.39, parameters - ILLUMINACLIP:2:30:10 SLIDINGWINDOW:4:20 MINLEN:25) (Bolger et al., 2014). SortMERNA (Kopylova et al., 2012) was used to remove any reads matching rRNA sequences. Remaining reads were aligned to the *C. elegans* reference genome (WBCel235) using HISAT2 (version 2.1.0, default parameters) (Kim et al., 2015). Read alignments were then counted using HTSeq-count (Anders, Pyl, & Huber, 2015) and gene counts inputted into DESeq2 (Love et al., 2014). RNA-Seq raw data have been deposited in the Gene Expression Omnibus (GEO) database under the accession number GSE146256.

## ACKNOWLEDGEMENTS

We would like to thank the Gurdon Institute Media Kitchen for their support providing reagents and media. We thank Kay Harnish for his support managing the Gurdon Institute Sequencing Facility. We thank Julie Ahringer’s lab for kindly sharing RNAi bacterial clones. We thank the National Bioresource Project (Tokyo, Japan) for providing the *nsun-5* deletion allele. We thank Claudia Flandoli for illustrating our working model. We are grateful for the Miska Laboratory members, especially Kin Man Suen, Eyal Maori and Grégoire Vernaz, for helpful discussions and advice, and Marc Ridyard for laboratory management and maintenance of our nematode collection.

---

**Supplementary Figure | 1 Referent to Figure 1.**
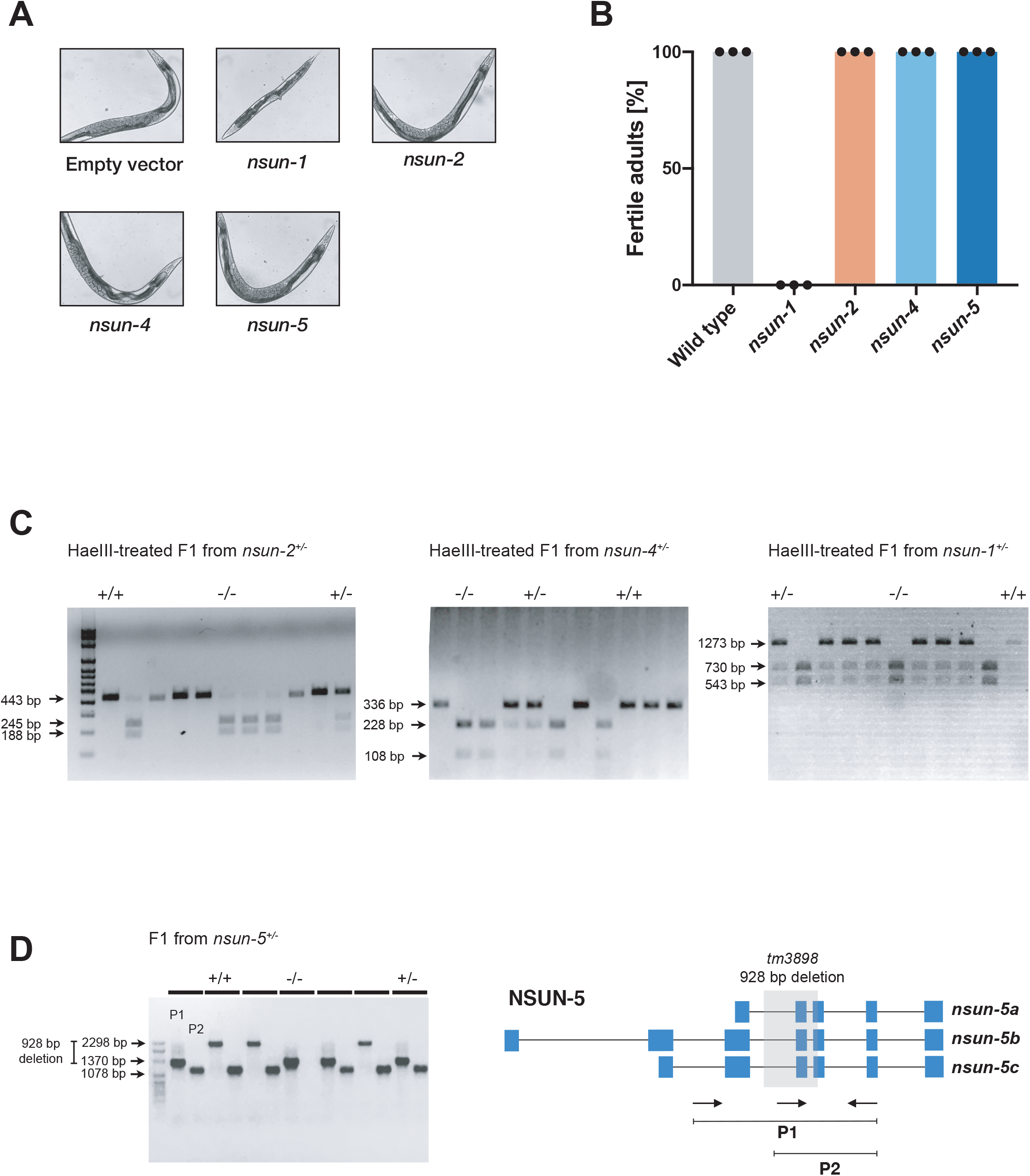
Knockdown and mutation of *nsun* genes. **(A, B)** Knockdown of *nsun* genes through RNAi by feeding. **(A)** Representative images of wild type adult animals after silencing of *nsun* genes via RNAi by feeding. Widefield DIC images are 10x magnification. **(B)** Percentage of fertile adults after gene silencing by RNAi. RNAi-treated animals were scored for fertility (presence of embryos in the germline) at the adult stage. Mean with SEM. 2 independent experiments, 3 biological replicates each. **(C)** HaeIII-treated DNA agarose gel showing genotypes of F1 individuals from heterozygous *nsun-1*, *nsun-2* and *nsun-4* mutants. **(D)** DNA agarose gel showing genotypes of F1 individuals from a heterozygous *nsun-5* mutant; diagram showing primers used for genotyping of *nsun-5* mutation.

**Supplementary Figure 2 | Referent to Figure 2.**
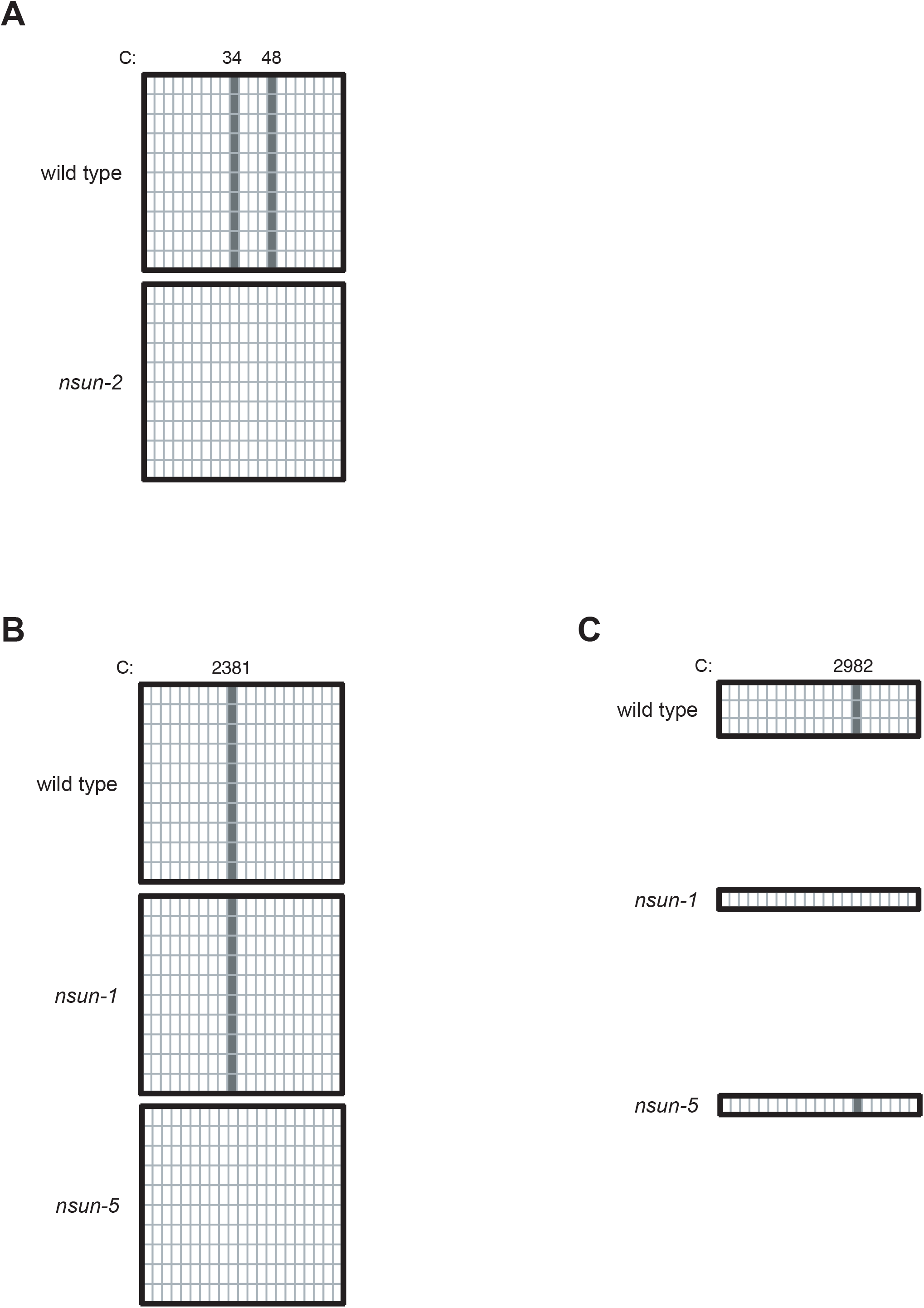
Enzymatic specificity of NSUN proteins in *C. elegans*. **(A, B, C)** Validation of enzymatic specificity of NSUN-2 (A), NSUN-5 (B) and NSUN-1 (C) by targeted bisulfite-sequencing. Each column represents one cytosine in the sequence of interest; each line represents one independent clone sequenced. Grey squares represent unconverted positions; white squares represent converted positions. 1 biological replicate.

**Supplementary Figure 3 | Referent to Figure 2.**
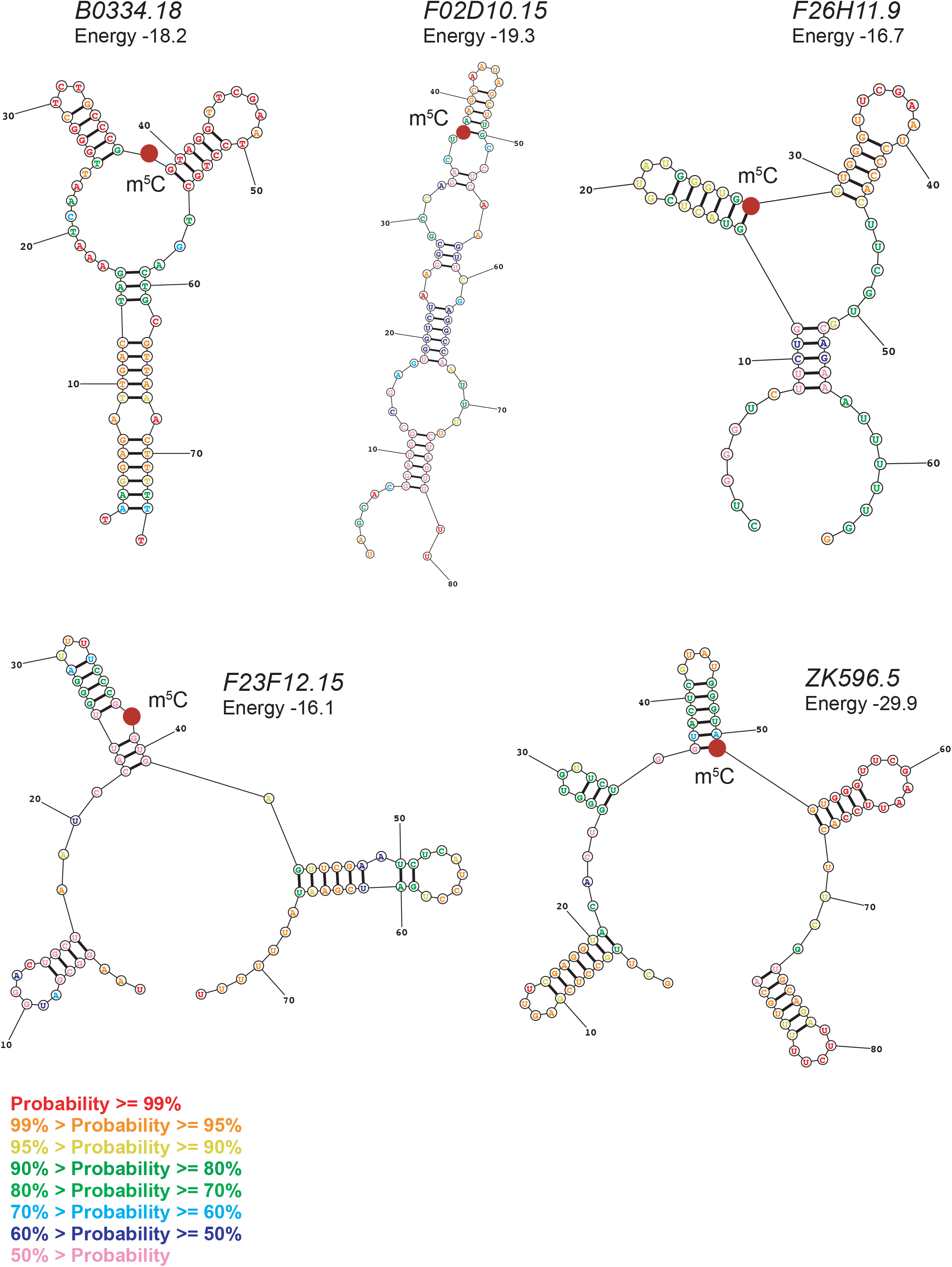
Predicted secondary structures of m^5^C methylated ncRNAs. Red dot indicates the methylated position. Structures predicted by the Predict a Secondary Structure Web Server (David Mathews Lab, University of Rochester) as the lowest free energy structures generated using default data.

**Supplementary Figure 4 | Referent to Figure 5.**
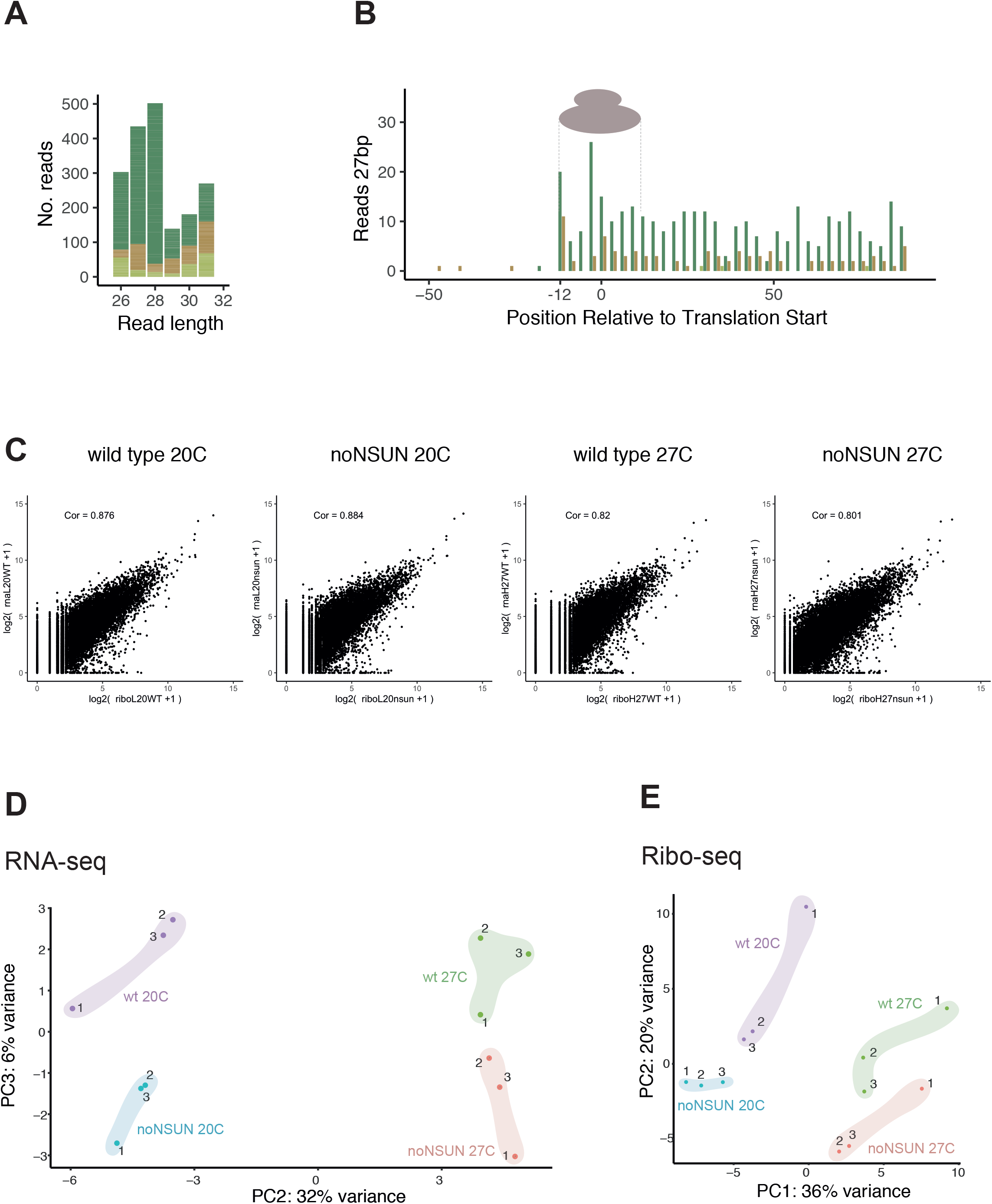
Quality assessment of ribosome profiling data. **(A)** Number of reads aligning the to the CDS in each frame after length stratification. The frame of the 5 prime nucleotide is shown. (**B)** Meta-gene example plot for reads that are 27 bp long showing the number of reads for each base pair of the start of each CDS. They are coloured according to the frame of their 5 prime base relative to the CDS. (**C)** Scatter plots showing the correlation between counts for RNA-seq and Ribo-seq for each gene in each sample. (**D)** PCA plot of RNA-seq counts for the 2000 genes with the highest variance. (**E)** PCA plot of Ribo-seq counts for the 2000 genes with the highest variance. 3 biological replicates.

**Supplementary Figure 5 | Referent to Figure 5.**
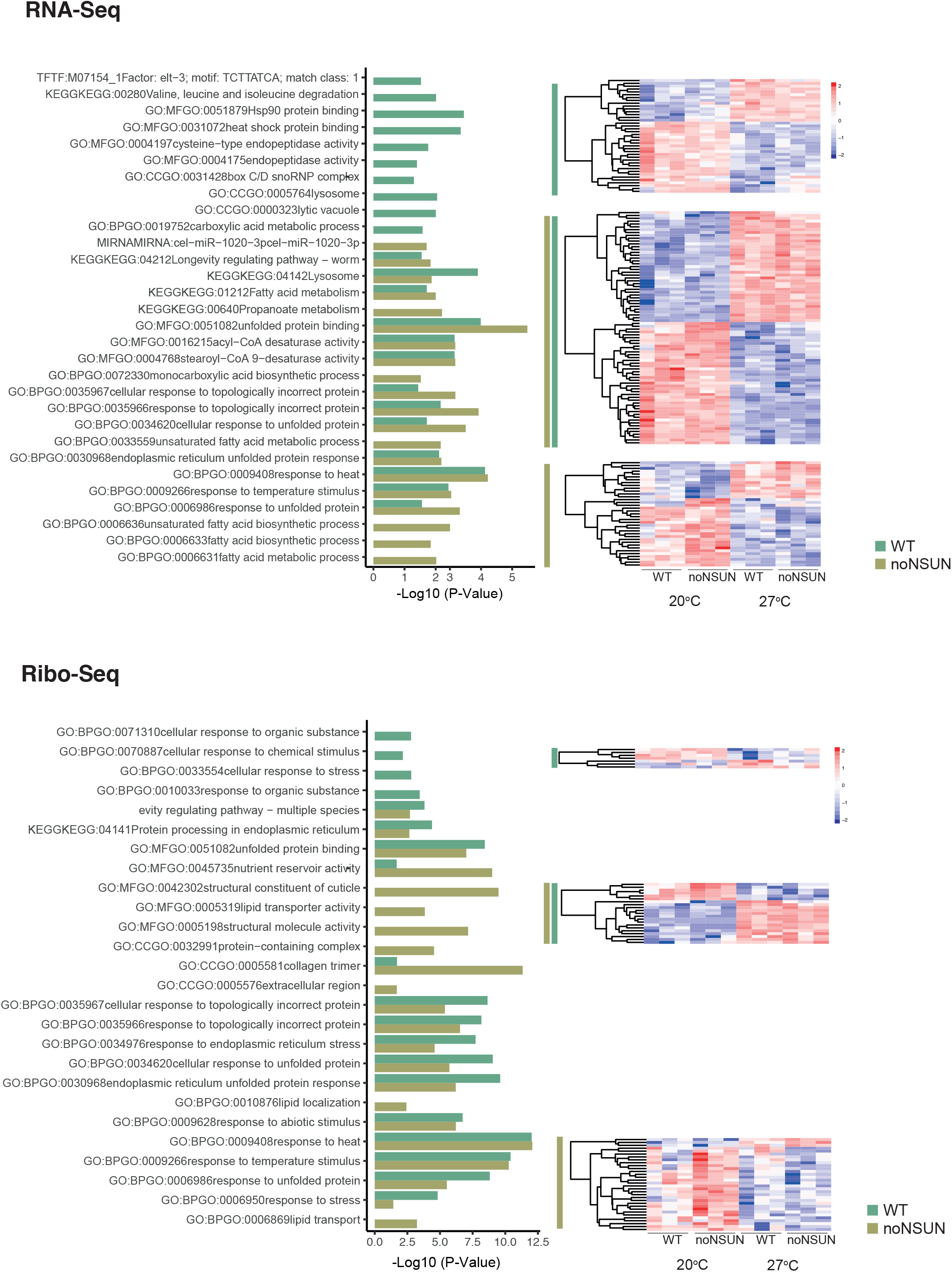
Differentially transcribed and translated genes upon heat shock. Heatmaps and gene ontology enrichment (biological process) analysis for the comparison between 20°C and 27°C in the wild type and noNSUN samples. Top shows RNA-seq (scaled normalised expression) and bottom shows Ribo-seq (scaled normalised RPFs). WT = wild type; 3 biological replicates.

**Supplementary Figure 6 | Referent to Figure 5.**
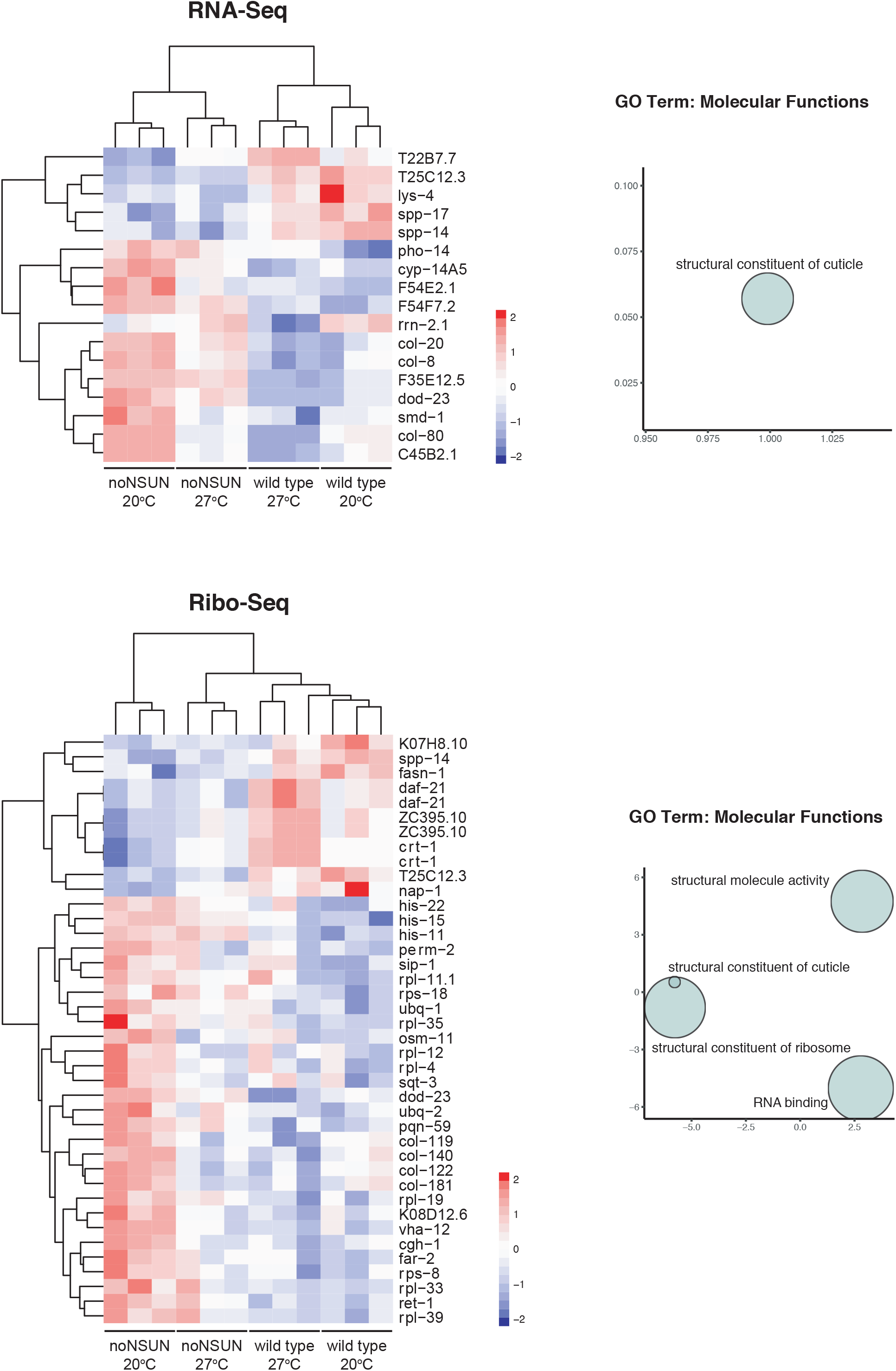
Differentially transcribed and translated genes upon loss of m^5^C. Heatmaps and gene ontology enrichment (biological process) analysis for the comparison between wild type and noNSUN samples. Top shows RNA-seq (scaled normalised expression) and bottom shows Ribo-seq (scaled normalised RPFs). WT = wild type; 3 biological replicates.

**Supplementary Figure 7 | Referent to Figure 5.**
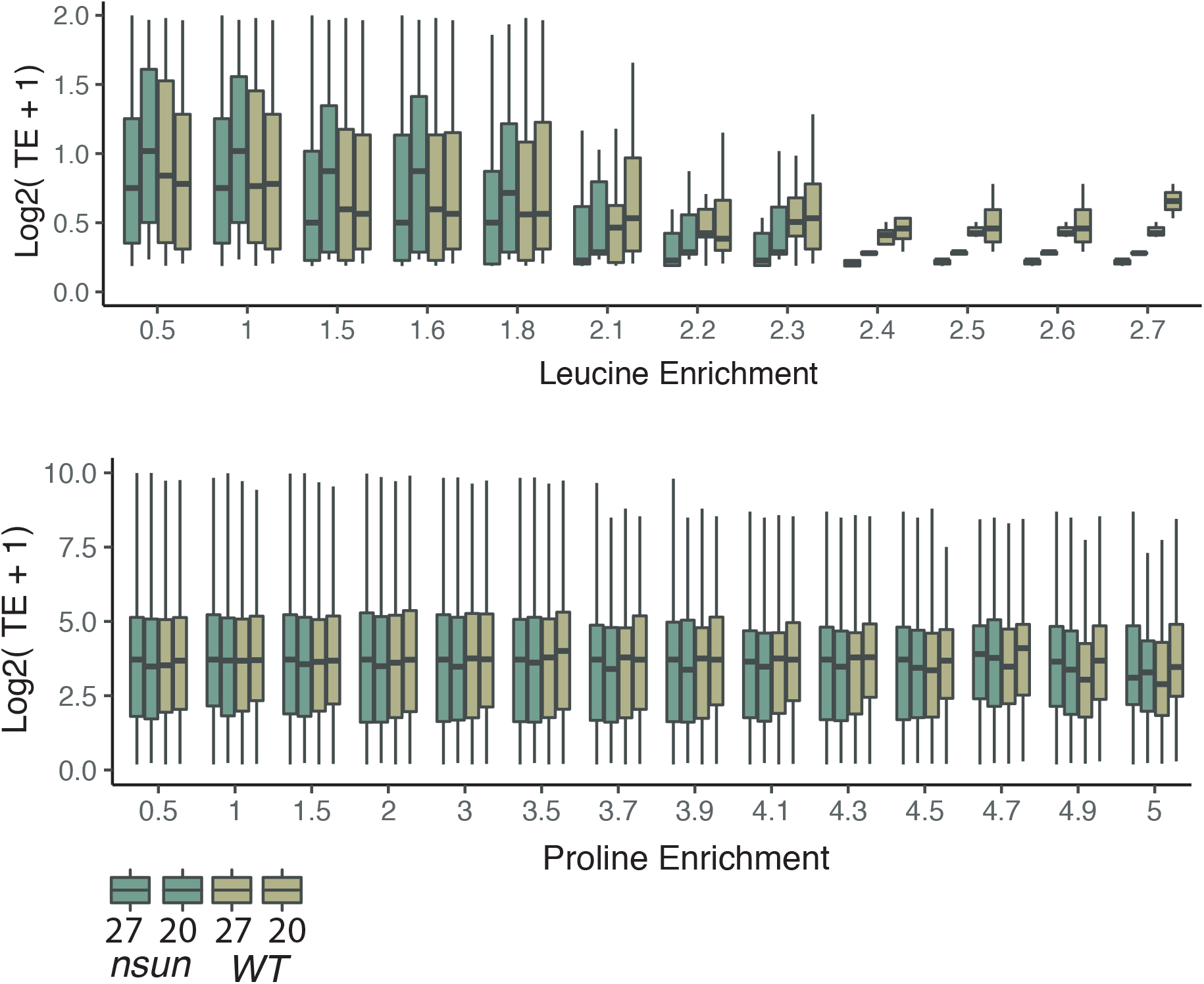
Translation efficiency of leucine and proline-enriched transcripts. Expanded from Figure 5C, translation efficiencies for genes with increasing enrichment for proline or leucine codons. 3 biological replicates.

